# ONECUT2 restricts Microfold cell numbers in the small intestine; a multi-omics study

**DOI:** 10.1101/2022.09.01.506202

**Authors:** Maria V. Luna Velez, Hannah K. Neikes, Rebecca R. Snabel, Yarah Quint, Chen Qian, Aniek Martens, Gert Jan C. Veenstra, Michael R. Freeman, Simon J. van Heeringen, Michiel Vermeulen

**Author notes:** These authors contributed equally to this work (Hannah K. Neikes, Rebecca R. Snabel). To whom correspondence should be addressed., Correspondence may also be addressed to: Maria V. Luna Velez. and, Correspondence may also be addressed to: Simon J. van Heeringen.

## Abstract

Microfold (M) cells reside in the intestinal epithelium of Peyer’s patches. Their unique ability to take up and transport antigens from the intestinal lumen to the underlying lymphoid tissue is key in the regulation of the gut-associated immune response. Here, we applied a (single-cell) multi-omics approach to investigate the molecular mechanisms that drive M cell differentiation in mouse small intestinal organoids. We generated a comprehensive profile of chromatin accessibility changes and transcription factor dynamics during *in vitro* M cell differentiation, allowing us to uncover numerous cell type-specific regulatory elements and associated transcription factors. Single-cell RNA sequencing resulted in the identification of an M cell precursor population. Our new computational tool SCEPIA determined that these precursor cells were characterized by high expression of and motif activity for the transcription factor ONECUT2. Subsequent perturbation experiments revealed that ONECUT2 acts downstream of the RANK/RANKL signalling to support Enterocyte differentiation and restrict M cell lineage specification *in vitro* and *in vivo*, thereby regulating mucosal immunity. This study provides a useful blueprint for future investigations of cell fate switches in the intestinal epithelium.

## Background

The Peyer’s patches (PP) are focal points within the small intestine, constituted by an organized lymphoid follicle and an overlying follicle-associated epithelium (FAE) (1,2). The FAE is a one-cell-thick layer composed mainly of Enterocytes and specialized epithelial cells called Microfold or M cells (1). M cells are important for enteral uptake and transport of various commensal and pathogenic microorganisms across the epithelial cell layer and to the subepithelial dome (3–6), where immune cells take on a defensive or tolerogenic response (1,7). Like all other cell lineages in the small intestine, M cells derive from intestinal stem cells (ISC) residing in the bottom of the intestinal crypts (8), however only a small percentage of ISC commit to the M cell lineage, making M cells one of the least abundant cell types in the small intestine (9,10). The polarization of M cells to PP and their low abundance reflect tightly regulated mechanisms restricting the commitment of progenitor cells into M cells, but it is precisely the low number of M cells that has made difficult the study of these mechanisms *in vivo*.

Intestinal organoids or “miniguts” are *in vitro* models capable of self-organizing into epithelial structures that molecularly, phenotypically and functionally resemble *in vivo* intestinal tissue (9,11). Minor modifications of the intestinal organoid culture medium can generate cell type-enriched organoids that can be used to decipher the molecular mechanisms that drive ISC fate (12,13). The receptor activator of nuclear factor κB (RANK) and its ligand (RANKL) were originally described for their key role in the induction of M cell differentiation (14,15). The addition of recombinant RANKL to the conventional intestinal organoid culture medium induces the differentiation of functional M cells *in vitro* (8), allowing the study of the fundamental signalling regulating M cell lineage specification.

In this study, we used a multi-omics framework on RANKL-treated mouse small intestinal organoids to define the gene-regulatory mechanisms underlying M cell differentiation. Changes in the morphology of organoids upon RANKL treatment were correlated to differences in protein levels, gene expression, histone modification and chromatin accessibility. By integrating bulk and single-cell RNA-sequencing data we were able to untangle complex gene-regulatory networks contributing to the M cell lineage specification in the intestine. Our findings led to the identification of an M cell precursor cluster, where the one cut domain family member 2 (ONECUT2) acts as a negative regulator of M cell lineage commitment in organoids and in the mouse small intestine.

## Results

### Generation of M cell-enriched mouse small intestinal organoids

*In vivo*, RANKL is expressed by MAdCAM-1^−^ mesenchymal cells beneath the FAE, functioning as M cell inducers by directly interacting with RANK-expressing progenitor epithelial cells (16,17). To mimic this mechanism *in vitro*, organoid culture medium was supplemented with recombinant RANKL for 6 days. Upon treatment, organoids acquired a cyst-like phenotype with wide, expanded villi regions and expression of glycoprotein 2 (GP2) in the apical membrane of a few cells (Fig. 1a and b). GP2 is an endocytic receptor for microorganisms and a marker for mature M cells (9,18). Our mouse small intestinal organoids carry a GFP reporter cassette at the locus of the leucine-rich repeat G-protein-coupled receptor (*Lgr5*) gene (19), a marker for ISC (20), which allowed us to visualize LGR5^+^ ISC in our organoid cultures. Concomitant to the increase in GP2 signal, RANKL treatment led to a reduction of LGR5 expression when compared to control organoids growing in conventional organoid culture medium (Fig. 1b). To evaluate the efficiency of M cell enrichment, flow-cytometry analysis was performed. In control conditions, the number of ISC and M cells was roughly 20% and <0.1%, respectively (Fig. 1c). Treatment with RANKL achieved a 4x increase in the number of M cells consistent with what is found *in vivo* (9,21) and an almost 50x decrease of ISCs (Fig. 1c).

**Fig. 1.**
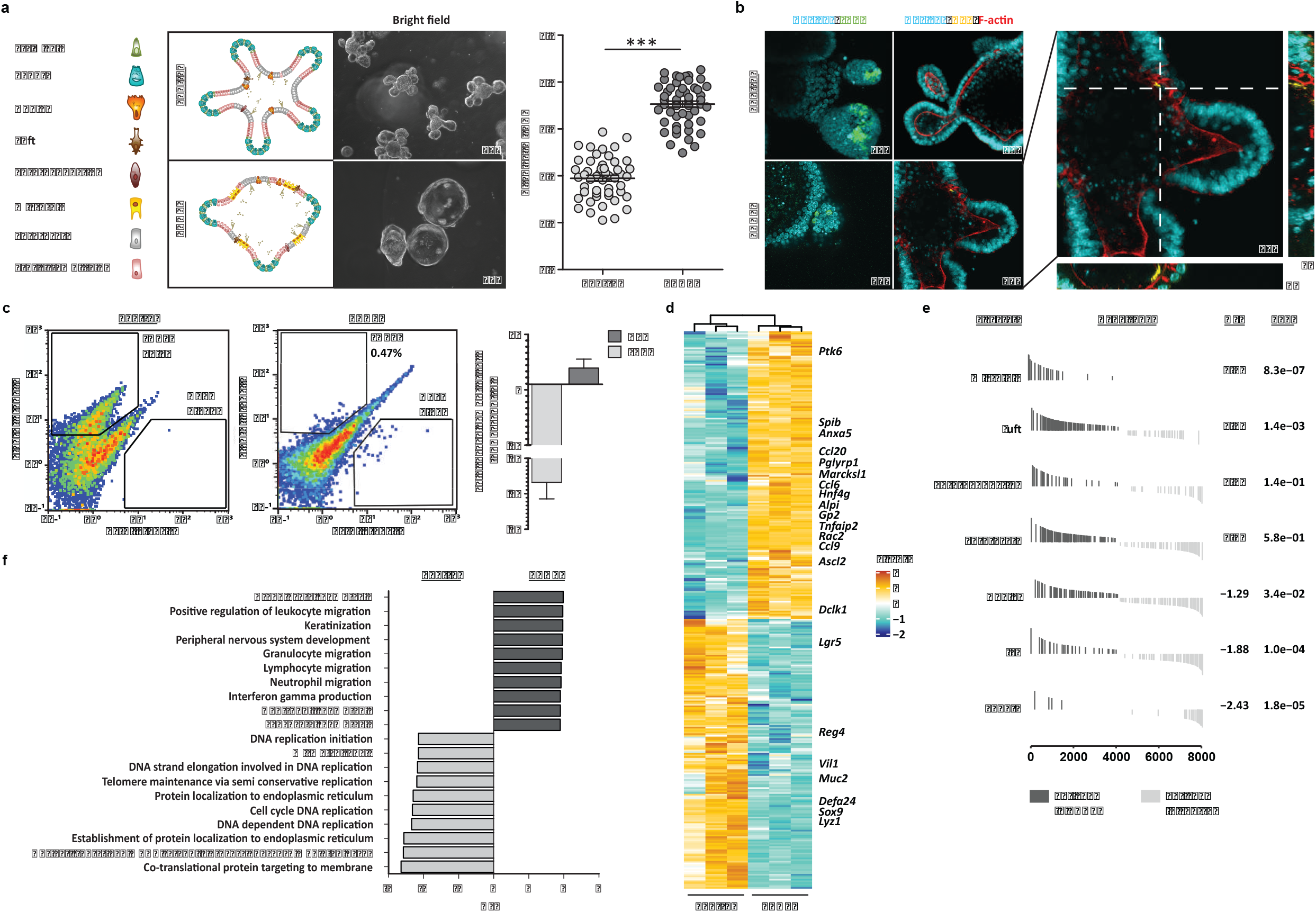
Generation of M cell-enriched mouse small intestinal organoids, **a** Brightfield images of control and M cell-enriched (RANKL-treated) organoids, next to graphical representation of their cell type composition. Microscope magnification is depicted. Graph shows the quantification of organoid circularity (AU, arbitrary units) and mean ± SEM. Control n=50 and RANKL n=51. **b** Nucleus (Hoechst), Factin (Phalloidin) and GP2 staining and detection of LGR5 signal in control and RANKL-treated organoids. On the right, orthogonal view on a zoom in image from the RANKL condition is displayed. Dotted lines show axes intersection point. Microscope magnification is depicted, **c** Representative density plots depict fluorescent intensities in the FITC (LGR5) and PE (GP2) channels, detected by flow cytometry. The percentage for each gate population is described. Bar graph represents the mean ± SEM of the fold change from at least five differentiation experiments. **d** Heatmap showing the relative change in mRNA expression upon RANKL treatment compared to control. Known markers of M cells and other intestinal cell lineages are highlighted. Rows show Z scores of normalized, log_2_-transformed values from significantly changing genes (padj < 0.05). **e** Gene-set enrichment analysis showing *in vivo* gene signatures of mouse small intestine cell lineages from Haber *et al*. (9) enriched in RANKL-treated organoids compared to control organoids (dark grey). Normalized enrichment score (NES) and padj values are described. Signatures enriched in control compared to RANKL condition (light grey) have a negative NES. f Gene-set enrichment analysis showing gene ontology (GO) terms significantly enriched (padj < 0.05) in control (light grey) and RANKL-treated organoids (dark grey).

In line with these findings, other known markers of M cells such as *Anxa5, Ccl20, Pglyrp1, Marcksl1, Ccl6, Gp2, Tnfaip2, Rac2* and *Ccl9* (9,18,22) were significantly upregulated in the RANKL conditions at a transcriptome level, while markers of ISC and other intestinal cell types showed opposite dynamics (Fig. 1d). To evaluate whether our *in vitro* model recapitulated *in vivo* tissue, whole-transcriptome changes induced upon RANKL-treatment were compared to cell type-specific intestinal signatures generated from a recently published single-cell survey of the mouse small intestine (9). The M cell and the Tuft cell gene signatures were significantly enriched under RANKL conditions, while the Paneth cell, ISC and Goblet cell signatures were enriched in control conditions (Fig. 1e). In addition, the M cell-enriched organoids were significantly enriched for genes involved in immune cell regulation but depleted in those associated with cell cycle and DNA replication (Fig. 1f), reflecting the physiological role of M cells. Together, these findings support the use of our *in vitro* model for the study of intestinal M cell differentiation with the potential of generating data of *in vivo* relevance.

### RANKL-induced transcriptional regulation

A significant dynamic transcript expression in 314 transcription factors was observed upon RANKL treatment in mouse small intestinal organoids (padj < 0.05). A fold change cut-off of >= 1 identified 45 and 55 transcription factors significantly upregulated in RANKL-treated and control organoids, respectively. Among those enriched in RANKL conditions we found *Relb, Spib* and *Sox8*, known drivers of M cell differentiation (23–25) (Fig. 2a). A prerequisite for transcription factor-mediated activity is an accessible chromatin, which is characterized by the presence of the H3K27ac modification, which marks active enhancers and, to a lesser extent, promoters (26). We used ATAC-sequencing and ChIPmentation to profile chromatin accessibility and the H3K27ac modification in the control and RANKL-treated organoid cultures. A total of 55,055 accessible sites were detected in the RANKL and control sequencing datasets, which were used to identify genomic locations that undergo a significant change in H3K27ac levels between both conditions. At these sites, we identified 93 and 123 motifs enriched in the RANKL and control conditions, respectively, further supporting an active regulatory role for 216 differentially expressed transcription factors (Fig. 2b).

**Fig. 2.**
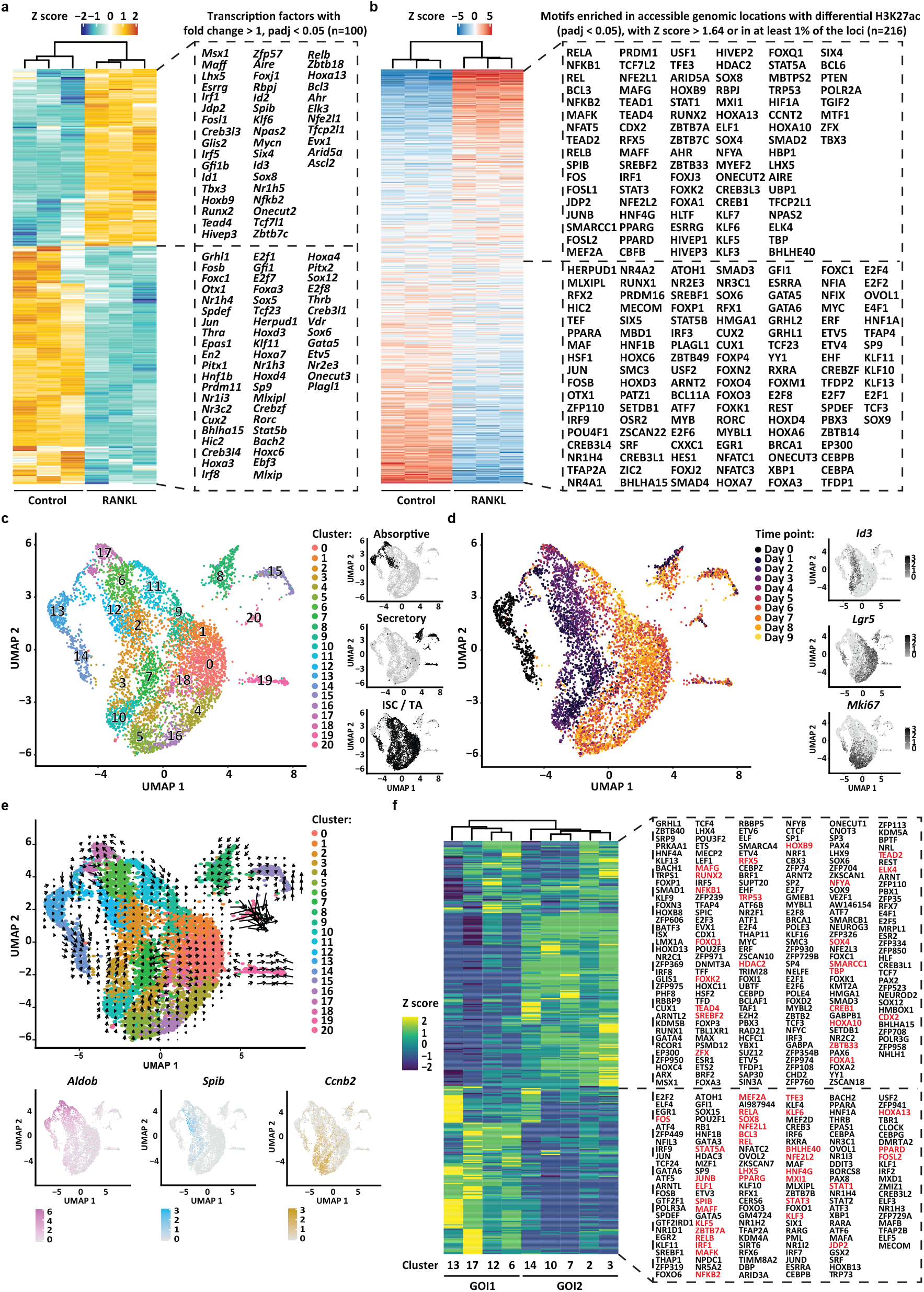
RANKL-induced transcriptional regulation. **a** Heatmap showing the relative change in mRNA expression of transcription factors upon RANKL treatment compared to control. Known markers of M cells and other intestinal cell lineages are highlighted. Rows show Z scores of normalized, log_2_-transformed values from genes significantly changing (padj < 0.05) with a fold change cut-off >1 and <-1. **b** Heatmap displaying enrichment of transcription factor motifs detected at accessible genomic locations (ATAC-sequencing data) that display significant dynamics of histone modification H3K27ac (ChIPmentation data, padj < 0.05) between RANKL-treated and control organoids. Only motifs of transcription factors differentially expressed (padj < 0.05) between RANKL-treated and control organoids from the bulk RNA-sequencing data are included. Rows show Z scores of motif enrichment with a cut-off of >1.64 and <-1.64. **c** UMAP embedding of single cell transcriptome from RANKL-treated organoids. Each dot represents a single cell. Cell colours represent cluster identity. Next, highlighted on black the partitions corresponding to cell populations expressing markers of the absorptive or secretory lineages, or markers of intestinal stem cells (ISC) and transit-amplifying cells (TA). **d** UMAP embedding overlay showing the harvesting time points during differentiation. Next, normalized and natural log-transformed expression of *Id3, Lgr5* and *Mki67*. **e** Velocity projections on UMAP embedding coloured as in **c.** Below, normalized and natural log-transformed expression of top markers from cluster 17 (*Aldob*), cluster 12 (*Spib*) and cluster 3 (*Ccnb2*). **f** Heatmap showing row Z scores of regulon activity (area under the curve scores) averaged per each cell population as analysed by pySCENIC. On red, regulons of transcription factors for which an active motif was found in b. Group of interest 1 (GOI1) and 2 (GOI2) are shown.

To obtain cell type-specific transcriptional regulation profiles across time, single-cell RNA-sequencing analysis was performed in RANKL-treated organoids harvested at different time points during treatment. In total, 6,636 cells were grouped into 20 clusters, which expressed markers from the absorptive or secretory lineages, or markers distinctive of ISC and daughter transit-amplifying cells (TA) (Fig. 2c, Additional file 1: Fig. S1a and Additional file 2: Table S1). Time seemed to separate cell clusters into two main partitions (Fig. 2d). One partition, with cells generated within the first five days of treatment, included cells expressing absorptive lineage makers (cluster 12, 13 and 17), a cell cluster expressing known M cell markers such as *Anxa5, Ccl20, Pglyrp1, Rac2, Serpinb1a* and *Ccl9* (cluster 12) and clusters expressing *Id3*, a marker for TA cells (cluster 2, 3, 7 and 10). A second time partition was constituted by cells originating from day 6 to day 9 (experimental end point), marked by the recovery of *Lgr5* expression and an increase in the number of secretory cells (Fig. 2d). *Mki67*, a marker of cell proliferation, seemed to be evenly distributed between cells from both partitions. In agreement with this, EdU labelling on RANKL-treated organoids showed that day 3 M cell-enriched organoids displayed a high number of LGR5^−^, proliferative cells (EdU-labelled), whereas day 6 organoids had proliferative and LGR5^+^cells solely at crypt bottoms (Additional file 1: Fig. S1b).

This time-resolved single-cell transcriptome data was suitable for RNA velocity calculations (27), an analysis in which spliced and unspliced RNA abundance is used to infer a ‘direction of change’ and predict the future state for each cell in a two-dimensional space. With the aim of unravelling the mechanisms driving M cell differentiation, we further narrowed analyses to cell clusters generated within the first five days of RANKL treatment. RNA velocity results suggested that day 0 *Lgr5^+^/Mki67^+^* cells (cluster 14) generated *Lgr5^−^/Mki67^+^* cells characterised by high *Ccnb2* expression (cluster 3) and that these cells turned in time into non-proliferative *Id3^+^* cells (*i.e*. cluster 2) (Fig. 2e, Additional file 2: Table S1). This process resembled the generation of TA cells from ISC observed *in vivo* (28). Interestingly, cluster 17, characterised by *Aldob* expression and originated from a second day 0 cluster (cluster 13), was shown as a precursor of *Spib^+^* M cells (cluster 12) (Fig. 2e, Additional file 2: Table S1). In agreement with these findings, clusters originating from cluster 13 (group of interest 1 or GOI1) displayed transcriptional regulatory units or ‘regulons’ (29) for the transcription factors NFKB2, RELB, SPIB and SOX8, well-established drivers of M cell differentiation. GOI1 also included regulons for HNF4G, ZBTB7B, RXRA and RARA, regulators of the absorptive lineage (12,13,30) (Fig. 2f). In contrast, group of interest 2 (GOI2) presented regulons of transcription factors previously associated with cell cycle control and crypt-villus maintenance such as MYC, E2F1 and E2F3 (31), and TEAD2, TEAD4 and SOX9 which regulate TA expansion (32,33). A total of 358 regulons supported the identity of cells from GOI1 and GOI2, from which a motif for 56 of these transcription factors was found within an accessible genomic region and significantly enriched by H3K27ac upon RANKL treatment (padj < 0.05) based on our bulk integrative analysis (Fig. 2f, highlighted in red). Collectively, our bulk and single-cell sequencing data defined the transcriptional and cellular responses to RANKL treatment in our organoid model and allowed the identification of a proposed M cell precursor cell population, cluster 17.

### ONECUT2 signalling in M cell precursor cell population

To predict transcription factor motif activity from single cell RNA-sequencing data, we designed a novel computational tool called SCEPIA (Single Cell Epigenome-based Inference of Activity) (34). By using computationally inferred epigenomes of single cells, SCEPIA can identify transcription factors that determine cellular states. In total, 81 transcription factors showed substantial motif activity within our single cell dataset, like *Pax4* and *Neurog3* in cells of Enteroendocrine cell identity (cluster 15), *Runx1* in Tuft cells (cluster 19), or *Spib* in M cells (cluster 12) (Additional file 2: Table S2). From those transcription factors highly expressed within GOI1 (cluster 6, 12, 13 and 17) *Hnf4g, Klf5, Onecut2* and *Tcf7l2* were the top four with the highest positive correlation between transcript expression and motif activity (Fig. 3a, Additional file 2: Table S2). The hepatocyte nuclear factor 4 gamma (HNF4G) is a key driver of Enterocyte cell differentiation (12,30). The krüppel-like factor 5 (KLF5) is a zinc-finger transcription factor expressed in both ISC and TA cells (35–37) while the one cut domain family member 2 (ONECUT2) transcription factor is decreased in LGR5^+^ ISC (38). The transcription factor 7-like 2 (TCF7L2) is a known downstream effector of the WNT pathway, expressed along the entire crypt-villus axis (39). All four transcription factors have been shown to be essential for proper differentiation of the embryonic intestinal epithelium (35,40–43), but their role in the modulation of gene expression during M cell differentiation was thus far unknown.

**Fig. 3.**
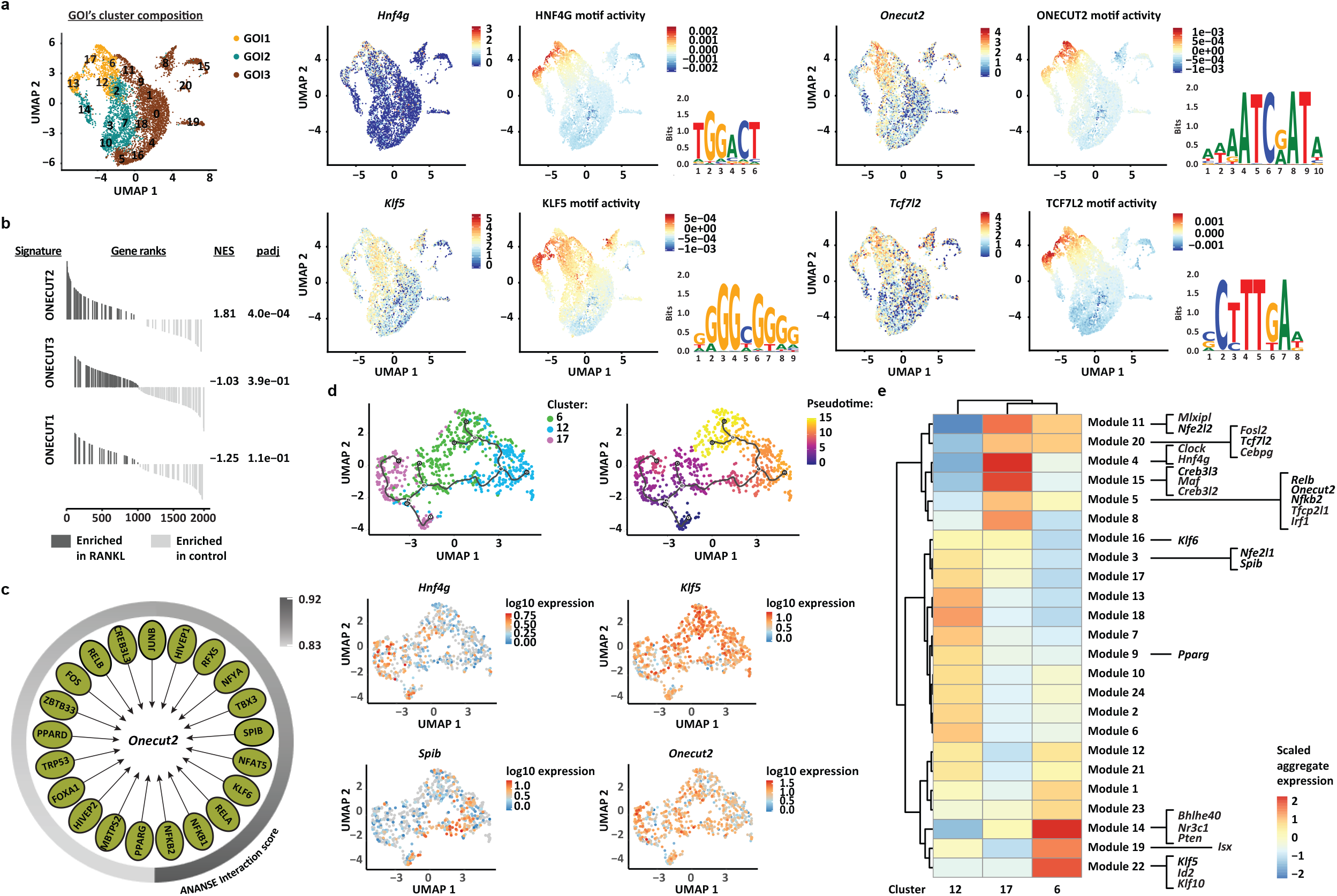
ONECUT2 signalling in M cell precursor cluster. **a** On the left, UMAP embedding overlay showing three partitions, group of interest 1 (GOI1), 2 (GOI2) and 3 (GOI3). On the right, top four transcription factors expressed within GOI1, with the highest positive correlation between transcript factor expression and motif activity. The logos for the motif of HNF4G (GM.5.0.Nuclear_receptor.0068), KLF5 (GM.5.0.C2H2_ZF.0006), ONECUT2 (GM.5.0.CUT_Homeodomain.0005) and TCF7L2 (GM.5.0.Mixed.0092) are displayed. **b** Gene-set enrichment analysis showing ONECUT1-, ONECUT2- and ONECUT3-specific gene signatures from Data ref: (48) enriched in RANKL-treated organoids compared to control organoids (dark grey). Normalized enrichment score (NES) and padj values are described. Signatures enriched in control compared to RANKL condition (light grey) have a negative NES. **c** Gene-regulatory network of *Onecut2* drivers predicted by ANANSE. Interaction scores (arbitrary cut-off ≥ 0.83) are depicted. **d** Trajectory plot overlaid with cluster identity of individual cells from RANKL-treated clusters in GOI1. Three major branches are shown along the plot. On the right, same trajectory plot with cells ordered along a pseudotime. Below, log_10_ expression values of *Onecut2* and top genes from each branching point are shown. **e** Co-regulated modules of differentially expressed genes by cells from RANKL-treated clusters in GOI1. The transcription factors predicted by ANANSE as *Onecut2* drivers and ONECUT2 targets, that are differentially expressed (FDR < 0.05) by cells from RANKL-treated clusters within GOI1 and defined as pseudotime-dependent genes are highlighted.

To define the context for HNF4G-, KLF5-, ONECUT2- and TCF7L2-mediated transcriptional regulation, a differential gene-regulatory network was built using ANANSE (44). Different to our integrative approach, ANANSE takes into account the transcription factor target gene expression in addition to transcription factor expression and H3K27ac and chromatin accessibility profiles to build a differential gene-regulatory network in RANKL over control conditions. A total of 15371 transcription factors and target genes constituted the gene-regulatory networks for HNF4G, KLF5, ONECUT2 and TCF7L2 (Additional file 2: Table S3). Interestingly, upon RANKL treatment, ONECUT2-binding motifs were found enriched in active regulatory elements from both *Tcf7l2* and *Klf5* (Additional file 2: Table S3), proposing ONECUT2 as an upstream regulator of TCF7L2 and KLF5-mediated transcriptional activity.

ONECUT2 forms part of the cut homeobox family of transcription factors and in mammals this family also includes ONECUT1 and ONECUT3 proteins (45–47). These three proteins have been shown to have a unique as well as an overlapping transcriptional output (48). To determine whether the enrichment for the ONECUT2-binding motifs determined by SCEPIA and ANANSE was specific for the ONECUT2 transcription factor, we performed a gene-set enrichment analysis (GSEA) with signatures unique for each cut homeobox protein. Only the ONECUT2 gene signature was shown to be significantly enriched upon RANKL treatment (Fig. 2b). Leading-edge (highly enriched) genes from the ONECUT2-specific signature encode proteins associated with the epithelial membrane, as well as transport and metabolism (Additional file 2: Table S4), in line with previous reports about the ONECUT2 signalling in the mouse small intestine (41). We, then performed ChIP-sequencing analysis using a ONECUT2-specific antibody. A total of 7261 ONECUT2 peaks were identified in the RANKL condition whereas binding of ONECUT2 was almost absent in control condition (Additional file 2: Table S5). Among others, upon RANKL treatment, ONECUT2 was found significantly bound (FDR < 0.05) to active chromatin regions upstream of *Tcf7l2, Klf5* and *Hnf4g* and of 167/364 ANANSE-predicted ONECUT2 target genes (Additional file 2: Table S5). This result places ONECUT2 as a master regulator in the GOI1. ANANSE also identified motifs for 21 transcription factors within *Onecut2* enhancer regions, among which those for RELB and NFKB2 binding, the latter having the highest predicted interaction score (Fig. 3c). Consistently, *Onecut2* was found part of the regulon of RELB (Additional file 2: Table S6). Heterodimer formation between RELB and the NFKB2 p52 subunit is known to be required for transcriptional activation of *Spib* and *Sox8* (22), proposing the involvement of the RELB-p52 dimer in the regulation of *Onecut2* expression.

To observe the process of M cell lineage commitment in more detail, we performed an unsupervised trajectory analysis in the RANKL-treated cell populations from the GOI1. Cells were ordered based on transcriptome expression and plotted as a function of pseudotime (Fig. 3d). This analysis showed that a subset of cells within precursor cluster 17, corresponding to day 0 (Additional file 1: Fig. S1c), gives rise to three cell populations, characterized by high expression of *Spib, Hnf4g* or *Klf5* (Fig. 3d). This is consistent with our experimental time points, where the *Spib^+^* cell population (M cells) was shown to differentiate earlier, followed by the *Hnf4g^+^* and later the *Klf5^+^* cell populations (Additional file 1: Fig. S1c). We, next investigated how the genes from the *Onecut2* regulatory network organized along the differentiation trajectories. For this, we included the ONECUT2 targets determined by ChIP sequencing and ANANSE-predicted *Onecut2* drivers, and only transcription factors that were differentially expressed within RANKL-treated cells clusters of the GOI1. Including *Onecut2*, the expression of 26 transcription factors was found to be pseudotime-dependent (Additional file 2: Table S7). To illustrate their contribution to each pseudotime branch, co-expressed genes were further grouped into modules that co-regulate across the cell clusters (Fig. 3e, Additional file 2: Table S7). *Spib, Nfe2l1* and *Pparg* were assigned to modules of genes expressed at high levels in cluster 12, while *Klf5, Id2, Klf10, Bhlhe40, Nr3c1, Pten* and *Isx* were part of modules of genes highly expressed in cluster 6. *Creb3l2, Creb3l3, Hnf4g* and other genes belonging to the ONECUT2 transcriptional output were assigned to cluster 17 modules. Remarkably, *Nfkb2, Relb* and *Onecut2* shared the same expression module (Additional file 2: Table S7), with genes in common upregulated in cluster 17, supporting the proposed regulatory mechanism of *Onecut2* expression by the RELB/p52 axis. Collectively, these analyses showed that precursor cells within cluster 17 give rise to cluster 6, 12 and a second subcluster of 17 which based on their transcriptome profile fit with a TA, M cell and Enterocyte identity, respectively.

### ONECUT2 restricts M cell lineage specification

To investigate the functional consequences of the ONECUT2 protein on M cell differentiation, organoid cultures were treated with the ONECUT2 inhibitor CSRM617. This compound was designed to specifically bind to the ONECUT2 homeodomain (HOX), thereby inhibiting DNA binding and ONECUT2-mediated transcriptional activation (49). CSRM617 alone had little, if any, effect on the morphology of organoids when compared to control conditions (Fig. 4a). In contrast, addition of CSRM617 to RANKL medium greatly affected the phenotype previously observed with RANKL treatment, resulting in small-lumen organoids with angular crypts (Fig. 4a). Consistent with the observed phenotype, CSRM617 alone did not affect much the transcriptome profile, compared to control conditions (Additional file 1: Fig. S2a). Co-treatment with RANKL and CSRM617 compared to RANKL alone led to a significant downregulation (padj < 0.05) of 4317 genes, which included 1150 ONECUT2 target genes (Fig. 4b). Together, these results indicate that ONECUT2 signalling in mouse small intestinal organoids is RANKL-dependent.

**Fig. 4.**
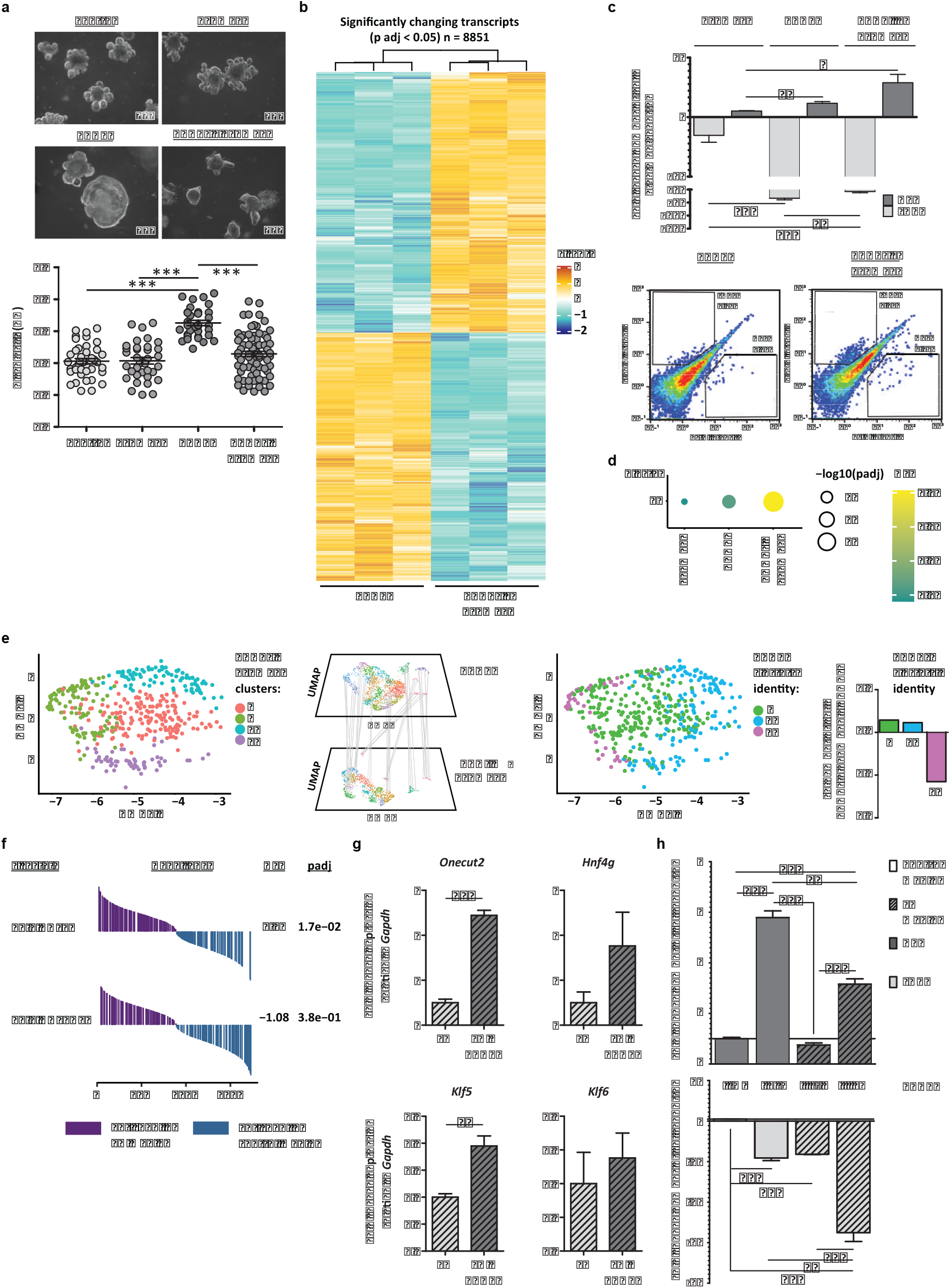
ONECUT2 restricts M cell lineage specification *in vitro*. **a** Bright-field images of organoids treated with RANKL, ONECUT2 inhibitor CSRM617 or a combination of both, and control organoids. Microscope magnification is depicted. Below, graph with organoid circularity quantification (AU, arbitrary units) and mean ± SEM. Control n=40, CSRM617 n=35, RANKL n=33 and RANKL + CSRM617 = 70. b Heatmap showing the relative change in the mRNA expression of transcripts upon co-treatment with RANKL and CSRM617, compared to RANKL-alone. Rows show Z scores of normalized, log_2_-transformed values from significantly changing genes (padj < 0.05). **c** Bar graph representing the mean ± SEM of at least four differentiation experiments. Below, density plots depicting fluorescent intensities in FITC (LGR5) and PE (GP2) channels, determined by flow cytometry. Percentage of gated populations is described. **d** Geneset enrichment analysis (GSEA) showing RANKL-treated cluster 12-specific gene signature, enriched in organoids treated as in **a,** compared to control. Dotplot shows the normalized enrichment score (NES, colour) and FDR (size). **e** Label transferring from RANKL-treated cell clusters within GOI1 to cells cotreated with RANKL and CSRM617. Bar graph on the right shows the fold change (FC) number of cells with RANKL clusters’ identity (RANKL + CSRM617 over RANKL). **f** GSEA showing the HNF4G- and ONECUT-ChIP signatures, enriched in organoids growing in EN medium, compared to control medium. NES and padj values are described. **e** Relative mRNA expression of *Onecut2, Hnf4g, Klf5* and *Klf6* detected in organoids growing on EN medium alone or with RANKL. **f** Ratio of LGR5^+^ or GP2^+^ cell numbers in organoids growing on EN or control medium with RANKL or EN medium alone, over control conditions, determined by flow cytometry. Bar graph representing the mean ± SEM of three differentiation experiments. For **b, c, e** and **f,** unpaired t test p < 0.05 is represented by (*), p < 0.01 by (**) and p < 0.001 by (***).

Given the fact that *Onecut2* was shown as a master regulator of GOI1, we hypothesized that inhibition of ONECUT2 genomic functions would have a negative effect on M cell differentiation. Strikingly, however, co-treatment with CSRM617 and RANKL resulted in an increase in the number of GP2^+^ cells (Fig. 4c). Effectively, co-treatment with CSRM617 resulted in a significant enrichment of the overall M cell signature from our single-cell dataset (Fig. 4d). These results indicated that ONECUT2 exerts an inhibitory effect in RANKL-induced M cell differentiation *in vitro*.

We, next assessed the transcriptome profile of RANKL-treated cell populations from GOI1 in single cells from organoids co-treated with RANKL and CSRM617 (Additional file 1: Fig. S2b). The transcriptome profile of precursor cluster 17 was almost completely lost in the co-treatment single-cell dataset (Fig. 4e), possibly reflecting an absolute induction toward the M cell lineage and supporting the role of ONECUT2 in the maintenance of cluster 17. The co-treatment also resulted in the loss of 147 regulons that were previously identified in RANKL-alone conditions, 48 of them corresponded to ONECUT2 target genes (Additional file 1: Fig. S2c, highlighted on red). It also included transcription factors characteristic of GOI2, a group of cell clusters largely composed by *Mki67^+^7Lgr5^−^* proliferative cells (Fig. 2d and f, Additional file 1: Fig. S2c). This is in agreement with the loss of the cystic phenotype that was observed in organoids subjected to RANKL treatment (Fig. 4a) and stresses a switch in cellular programs, from a balanced proliferative/differentiated state to an overdriven differentiation program.

If the induction of M cell differentiation by RANKL and CSRM617 co-treatment inhibited that of Enterocyte cluster 17 cells we wondered whether the induction of Enterocyte cell differentiation would then inhibit M cell number. In a previous study, we established the culture medium conditions for the enrichment of Enterocytes in mouse small intestinal organoids, the so-called EN medium (12). Interestingly, the HNF4G but not ONECUT2 signature was found significantly upregulated in EN conditions (Fig. 4f). Remarkably, the addition of RANKL to EN medium upregulated the levels of *Onecut2, Hnf4g*, and ONECUT2 targets *Klf5* and *Klf6*, and markedly lowered the number of M cells compared to regular RANKL medium (Fig. 4g and 4h). These experiments demonstrated that the ONECUT2 induction of Enterocyte differentiation and inhibition on M cell lineage specification *in vitro* is dependent on the RANK/RANKL signalling.

### ONECUT2 limits M cell number in Peyer’s patches

To investigate the physiological relevance of our findings *in vivo*, mice were treated with CSRM617 as described in Additional file 1: Fig. S2d. Because RANKL is naturally highly expressed within the Peyer’s patches (PP) of the small intestine (16,17), we assessed the protein expression of M cell markers in this tissue. Reassuringly, treatment with CSRM617 increased the number of SPIB^+^and GP2^+^cells in the PP, compared to control treatment (Fig. 5a). Consistently, treatment induced an upregulation of M cell markers *Spib, Gp2* and *Tnfaip2* in the small intestine tissue, all of which increased in time (Fig. 5b). While the expression of the transcription factor and M cell driver *Spib* was significantly downregulated immediately after termination of the treatment, mature M cell markers *Gp2* and *Tnfaip2* (9) remained expressed at levels higher than in control conditions even after a week of recovery from treatment, likely reflecting the turnover of M cells (Fig. 5a). Treatment achieved downregulation of ONECUT2-targeted genes *Klf5, Klf6, Hoxb8, Foxn3* and *Tfcp2l1*, similar to the effect observed in organoids (Fig. 5c). ONECUT2 inhibition also led to a reduction in the levels of *Hnf4g* (Fig. 5c), but did not affect the transcript levels of markers from other cell types in the small intestine such as *Dclk1* (Tuft), *Defa6* (Paneth) or *Reg4* (Enteroendocrine) (Additional file 1: Fig. S2e), reiterating on the RANKL-dependent role of ONECUT2 to cells of the absorptive lineage.

**Fig. 5.**
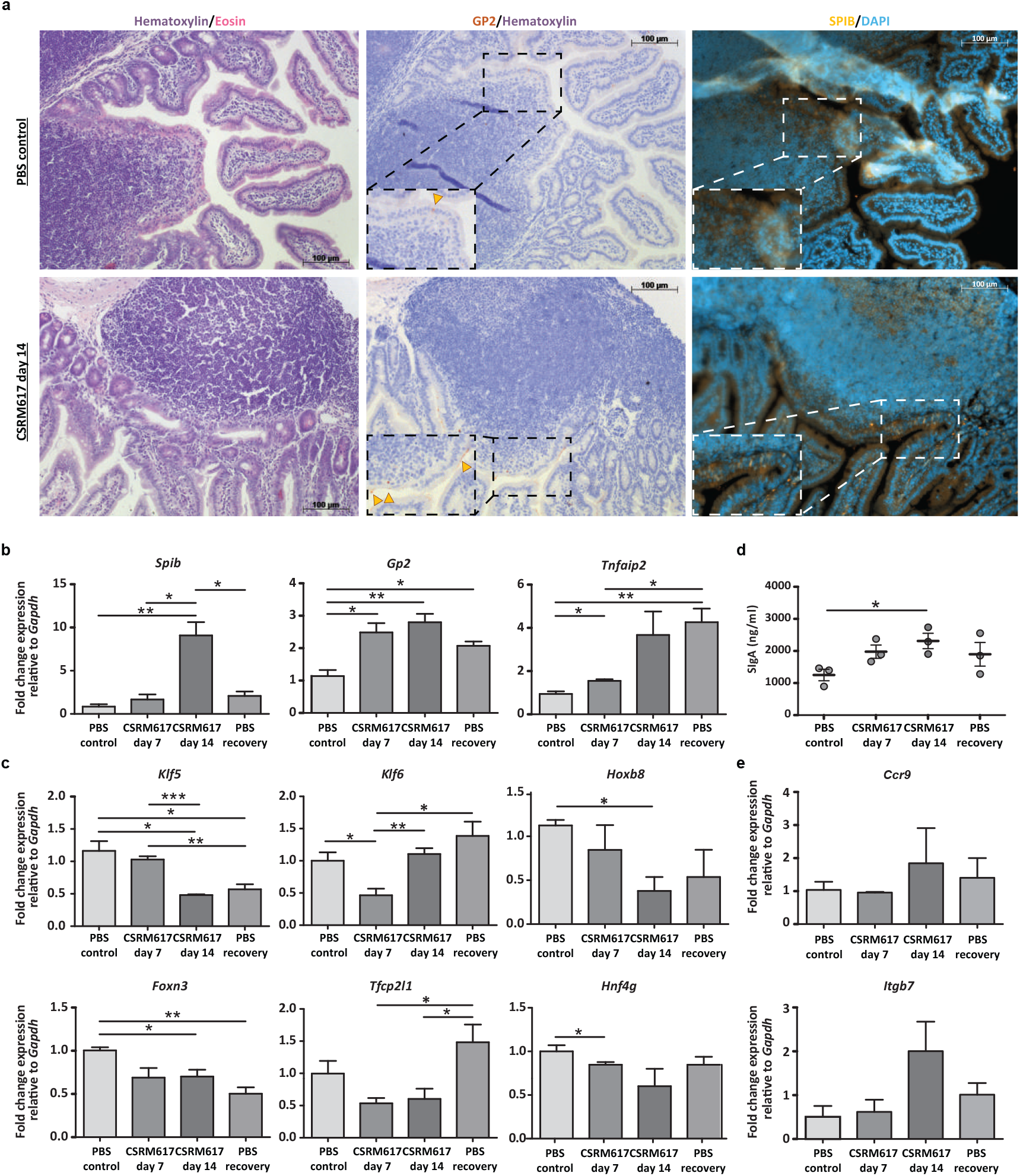
ONECUT2 regulates M cell number and mucosal immunity *in vivo*. **a** Immunohistochemistry analysis for GP2 and SPIB protein expression in Peyer’s patches of small intestine sections from mice treated with PBS control or CSRM617 for 14 days. Nuclear counterstaining with hematoxylin (bright field) or DAPI (fluorescence) is shown. Scale bars are annotated. Dotted rectangles show amplified areas and yellow arrow heads show GP2^+^ cells, b RT-qPCR analysis showing relative mRNA expression of M-cell markers *Spib, Gp2* and *Tnfaip2* in small intestine sections from mice treated as described in Additional file 1: Fig. S2d. Unpaired t test p < 0.05 is represented by (*) and p < 0.01 by (**). c Relative mRNA expression of ONECUT2-targeted genes *Klf5, Klf6, Hoxb8, Foxn3, Tfcp2l1* and *Hnf4g* detected in tissue from **b.** Unpaired t test p < 0.05 is represented by (*), p < 0.01 by (**) and p < 0.001 by (***). **d** Secretory IgA (SIgA) levels in faecal samples from mice treated as described in Additional file 1: Fig. S2d. Unpaired t test p < 0.05 is represented by (*). **e** Relative mRNA expression of markers from IgA^+^ B cells *Ccr9* and *Itgb7* detected in tissue from **b.**

The organized lymphoid follicle underlying the PP are sites for the generation of IgA^+^ B cells and the production of secretory IgA (SIgA) (7), the body’s first line of defence in the resistance against toxins and bacterial and viral infections (50). SIgA production is a consequence of M cell immune-related functions and its expression level reflects M cell numbers (15,51,52). Because ONECUT2 inhibition led to an increase in the number of M cells, we investigated to what extent this could affect the levels of SIgA. To this aim, we run ELISA experiments for targeted detection of mouse IgA in faecal samples of treated mice. Faecal SIgA levels increased over time after treatment with CSRM617 and dropped after a week receiving control treatment (Fig. 5d). These levels correlated with the expression of *Ccr9* and *Itgb7*, markers of IgA^+^ B cells (53) (Fig. 5e). The weight of mice treated with CSRM617 was comparable to that of littermate controls and no difference was observed between genders (Additional file 1: Fig. S2f). Based on these observations, we conclude that ONECUT2 restricts the M cell number in PP, indirectly regulating B cell functions and SIgA production.

## Discussion

In this study, we set out to use multiple omics platforms and diverse computational tools to define the molecular landscape of M cell-enriched organoids in an unbiased, comprehensive manner. By combining the strengths of bulk sequencing methods (*e.g*. small input material, sequencing depth) and single-cell approaches (*e.g*. characterisation of small populations, trajectory inference tools) to generate time-resolved data, we untangled complex gene-regulatory networks that contribute to the M cell lineage specification.

Our integrative approach revealed that ONECUT2 acts as a negative regulator of M cell differentiation *in vitro* and *in vivo*, by supporting Enterocyte differentiation in a RANKL-dependent manner. Under RANKL conditions, blockage of ONECUT2-mediated transcriptional activity led to an increase in the number of M cells (*Spib^+^, Anxa5^+^*) and a decrease in the Enterocytes cell population (*Hnf4g^+^, Alpi^+^*). A low M cell number has been proposed to be needed for the maintenance of proper intestinal barrier function and to prevent an over-reactive immune response (54), and ONECUT2 expression could be one of the mechanisms ensuring this. The elevated expression of IgA^+^ B cell markers and high levels of intestinal SIgA we observed *in vivo* upon ONECUT2 inhibition supports this hypothesis. Moreover, the expression of *Klf5*, a transcription factor predicted to be target of ONECUT2 has been shown to protect mice against murine colitis by activating the JAK-STAT signalling pathway (55).

*Onecut2* was one of the transcription factors with the highest expression and motif activity in a cell population precursor for M cells, Enterocytes and a TA cell population. Expression of *Onecut2* was correlated with that of *Nfkb2* and *Relb*, transcription factors responsible for transcriptional activation of *Spib* and *Sox8* (22) and which expression in the small intestine are restricted to follicle-associated crypts (56). Motifs from both transcription factors were found within *Onecut2* enhancers, supporting a potential role for RELB-p52 dimer in the transcriptional regulation of *Onecut2*.

We have created a new computational tool we named SCEPIA (34) and validated it for its capacity to predict transcription factor motif activity from single cell RNA-sequencing data, to identify those transcription factors, like ONECUT2, that are determinant of cellular states. Likewise, we have generated several resource datasets that can be used to perform different analyses, for instance, to uncover additional regulatory axes underlying RANKL signalling, or as reference datasets of the chromatin remodelling and transcriptional responses to ONECUT2-transcription factor activity. In human and mouse, ONECUT2 is highly expressed in the liver, pancreas, small intestine and the nervous system. In addition, ONECUT2 expression is significantly associated with poor clinical outcome in diverse type of cancers such as prostate, breast, gastric, colon, clear cell renal, brain, and lung cancer (49). The systemic administration of the ONECUT2 inhibitor CSRM617 in mice, described in this study, generated tissue resources that can be used to study the many diverse roles of the ONECUT2 in different tissues, in the context of health and disease.

### Limitations of study

Mouse small intestinal organoids are a powerful and versatile *in vitro* model, which resembles the *in vivo* intestinal epithelium at a cellular, molecular and functional level, thereby facilitating the study of complex biological processes such as stem cell renewal and cell differentiation. Though organoids typically consist of several distinct epithelial cell types, cultures rarely include other non-epithelial cells (*i.e*. stromal, mesenchymal or immune cells), which represent a source of factors that can be important for the cellular maturation and/or differentiation of epithelial cells. For M cells, beside MAdCAM-1^−^ mesenchymal cells, which express RANKL, lympho-epithelial cell interactions such as those with B cells have been shown to play a role in the induction of M cell differentiation (57,58). In addition, it has been shown that M cell differentiation can be induced in response to external stimuli such as a pathogen infection (56) but incorporating all these factors in a culture with organoids can be challenging. Thus, it remains to be answered whether pathogen-induced signalling would overrule inhibitory M cell differentiation signals such as those mediated by ONECUT2, uncovered in this study.

## Conclusions

In this study we have shown that the ONECUT2 transcription factor acts downstream of RANK/RANKL signalling to orchestrate Enterocyte and M cell lineage specification *in vitro* and *in vivo*. Our perturbation studies demonstrated that ONECUT2 in fact restricts M cell differentiation within the FAE which has an impact in mucosal immunity as the levels of SIgA, the body’s first line immune defence, is a direct consequence of M cell function.

While the focus of this study resulted in the discovery of a transcription factor as a regulator of intestinal M cell differentiation, the computational workflows described here can serve as an outline for future endeavours aimed at deciphering gene expression regulation of stem cell fate and differentiation in the intestine and other multicellular tissues. Over the last years, different computational tools have been designed to model complex gene-regulatory networks and to facilitate the integration of multiple data types *i.e*. transcriptome, genome) of different resolutions *i.e*. bulk, single-cell). Though each tool is informative on its own, we found that the integration of all these was essential for the elaboration of the conclusions presented in this study.

## Methods

### *In vivo* CSRM617 experiments

Both male and female C57BL/6 mice (5 weeks old) were purchased from Charles River. After arriving, the mice were quarantined for one week prior to the study. Four treatment groups were designed for the *in vivo* experiments with the ONECUT2 inhibitor CSRM617 (Additional file 1: Fig. S2d). Each treatment group contained 3 male and 3 female C57BL/6 mice of 6-6.5 weeks of age. Briefly, 6 mice received 50□mg□kg-1 CSRM617 (Cedars Sinai Medical Center) for 7 days and 6 mice for 14 days. A recovery group consisted of 6 mice treated with 50□mg□kg-1 CSRM617 (Cedars Sinai Medical Center) for 7 days followed by 7 days treatment with phosphate buffered saline (PBS) and 0.2% dimethyl sulfoxide (DMSO, Sigma-Aldrich). Lastly, a control group treated with PBS and 0.2% DMSO for 7 days was included. All treatments were performed by intraperitoneal injection. Body weight was tracked 3 times per week. For ELISA experiments, faecal samples were collected 30 minutes before sacrifice of the mice. A total of 100 mg of mice faeces were dissolved in 1 mL of 1x PBS with 1% protease inhibitors (Thermo Scientific) and vortexed for 15 minutes at 4°C. Supernatant was isolated after centrifugation at 12,000 x g for 10 min and stored at −80°C until IgA measurements. The small intestines of 3 mice per treatment group (1 female and 2 males) were gently rinsed twice with 1x PBS and collected in individual tubes containing 1.2 mL of RNAlater Stabilization Solution (Invitrogen), and frozen at −80°C until being processed for RNA isolation. The small intestines from the remaining 3 mice of each treatment group (2 female and 1 male) were collected to prepare separate formalin-fixed paraffin-embedded tissue blocks following the “swiss roll” method (59).

### Culture of mouse small intestinal organoids

The mouse small intestinal organoids used in the current study were generated by isolating LGR5+ adult stem cells from the gut of a female *Lgr5*^GFPDTR/+^ mouse and culturing them in a well-defined semi-solid culture medium supplemented with essential growth factors (11). The female *Lgr5*^GFPDTR/+^ mouse was generated by replacing the first coding exon of *Lgr5* with a cassette containing the coding sequence for the enhanced green fluorescent protein (EGFP) linked in frame to the human diphtheria toxin receptor (DTR) gene cDNA (19).

Small intestinal organoids derived from a female *Lgr5*^GFPDTR/+^ mouse (19) were cultured as described previously (11). Briefly, organoids were grown embedded in a mix of 90% ice-cold RGF BME Type 2 PathClear (Cultrex, R&D Systems) and 10% Dulbecco’s modified Eagle’s Medium/Ham’s F12 (Gibco), in a humidified atmosphere at 37°C and 5% CO2. For maintenance and control conditions organoids grew in ENR medium which contained advanced Dulbecco’s modified Eagles Medium/F12 (Gibco) supplemented with 1x Penicillin-Streptomycin (Gibco), 10 mM HEPES (Gibco), and 1x GlutaMAX Supplement (Gibco), 1x B27 (Gibco), 1.25 mM N-acetylcysteine (Sigma Aldrich), 50 ng/mL mEGF (Gibco), 10% final volume NOGGIN conditioned medium, 5% final volume R-SPONDIN1 conditioned medium. For maintenance, organoids were split by mechanical dissociation using a Pasteur pipette every 5 days and the culture medium was refreshed every other day (11). For microfold (M) cell enrichment, organoids were cultured in ENR supplemented with 3 μM CHIR99021 (Axon) and 1.5 mM valproic acid (Sigma) for three days, followed by one day in ENR medium. Afterwards, treatment with 100 ng/mL of recombinant mouse TRANCE (RANKL) (BioLegend) was done for 6 or 9 days, alone or in combination with 40 μM CSRM617 Hydrochloride (Sigma-Aldrich). For EN experiments, RANKL was added to EN medium described previously (12). ENR medium alone or in combination with 40 μM CSRM617 were used as controls. Medium was refreshed every three days.

### Bulk RNA-sequencing sample preparation

After treatment, organoids from three independent differentiation experiments were harvested using Organoid Harvesting Solution (Cultrex) according to manufacturer’s protocol. Pellets were snap frozen and stored at-80 °C until being processed for RNA isolation using the RNeasy RNA extraction kit (Qiagen) with *DnaseI* treatment using RNase-Free DNase (Qiagen). Total RNA (1 μg per replicate) was used for library preparations using the KAPA RiboErase Kit (HMR) (Roche) following manufacturer’s protocol. A fragmentation step was carried out for 6.5 minutes at 94°C and 7 μM NextFlex DNA barcodes (PerkinElmer) were used for adapter ligation. Libraries were amplified using 6 amplification cycles. Library concentration was measured using the dsDNA Fluorescence Quantification Assays (DeNovix), and library size was determined using the BioAnalyzer High Sensitivity DNA Kit (Agilent Technologies) in a BioAnalyzer instrument (Agilent). Sequencing was performed using an Illumina NextSeq 500, and 42-bp paired-end reads were generated.

### Bulk ATAC-sequencing sample preparation

After harvesting of organoids from three independent differentiation experiments, pellets were cryopreserved in Recovery Cell Culture Freezing Medium (Gibco) at −80°C until being processed for ATAC-sequencing (ATAC-seq). ATAC-seq libraries were prepared following an adapted protocol from Buenrostro *et al*. and Corces *et al*. (60,61). In brief, 50,000 cells per replicate were lysed in 50 μl of icecold ATAC Resuspension Buffer (RSB) containing 0.1%IGEPAL (Sigma), 0.1% Tween-20 (Sigma), and 0.01% Digitonin (Sigma) for 3 minutes on ice. Lysates were resuspended with 1 mL of cold ATAC-RSB containing 0.1% Tween-20 (Sigma) and nuclei were pelleted by centrifugation at 500 rcf for 10 minutes at 4°C in a fixed angle centrifuge. Nuclei were tagmented for 6 minutes at 37°C with in-house made *Tn5* enzyme while shaking at 650 rpm. The tagmentation reaction was stopped with stop buffer (44 mM EDTA, 131 mM NaCl, 0.3% SDS, and 600 μg/ml proteinase K as final concentration). Genomic DNA was purified using 2x AMPure XP beads (Beckman Coulter) and subsequently amplified by PCR using KAPA HiFi Hotstart Ready Mix and Nextera Index Kit (Illumina) primers. Lastly, a reverse-phase 0.65x AMPure XP beads (Beckman Coulter) DNA purification step was done, followed by 1.5x AMPure XP beads (Beckman Coulter) purification. Library concentration was measured using the dsDNA Fluorescence Quantification Assays (DeNovix), and library size was determined using the BioAnalyzer High Sensitivity DNA Kit (Agilent Technologies). DNA libraries were sequenced with an Illumina NextSeq 500 at a read length of 38 bp.

### ChIPmentation sample preparation

Following harvesting, organoids were crosslinked with 1 % formaldehyde in 1x PBS shaking at 650 rpm for 10 minutes at room temperature, and glycine (125 mM final concentration) was added to quench the reaction. After a washing step with 1x ice-cold PBS, cell pellets were snap frozen and stored at −80°C until being processed for ChIPmentation. A total of 250,000 cells per replicate were lysed in a buffer containing 20 mM HEPES pH 7.6, 1% SDS, 1x Protease Inhibitor Cocktails (Roche). Chromatin was sheared using the medium setting of the Biorupter Pico (Diagenode) with 6 cycles (20 seconds on/30 seconds off) to achieve fragments between 200 and 600 bp. Samples were then spun at 13,000 rpm for 5 minutes at room temperature and supernatant containing fragmented chromatin was used for ChIPmentation as described by Schmidl *et al*. (62) with the modifications published in Lindeboom *et al*. (12). Briefly, before starting the immunoprecipitation, we saved 1% of the input material of either control or RANKL-treated organoids and stored it at 4°C until further processing (tagmentation). Then, chromatin of each replicate was incubated with 1 μg of antibody against H3K27ac (Diagenode, C15410196, RRID:AB_2637079) or 1 μg of antibody against ONECUT2 (Proteintech, 21916-1-AP, AB_2848180) in dilution buffer (1% Triton X-100, 1.2 mM EDTA, 16.7 mM Tris pH 8, 167 mM NaCl, 1x Protease Inhibitor Cocktail (Roche)), rotating overnight at 4°C. Antibody-bound chromatin was purified using protein A and protein G Dynabeads (Invitrogen) followed by two washing steps with increasing concentration of NaCl and one washing step with TE buffer as described before (12). Tagmentation of immunoprecipitated and input chromatin was performed with in-house generated *Tn5* enzyme for 10 minutes at 37°C while shaking at 550 rpm, followed by washing steps with decreasing amounts of NaCl and soap (12). Lastly, samples were de-crosslinked with 20 μg of proteinase K (Sigma) in elution buffer (0.5% SDS, 300 mM NaCl, 5 mM EDTA, 10 mM Tris pH 8) for 1 hour at 55°C and 1,000 rpm shaking followed by an overnight incubation at 65°C and 1,000 rpm shaking. The eluted, tagmented chromatin was treated with 10 μg proteinase K (Sigma) for 1 additional hour at 55°C and purified using 2x AMPure XP beads (Beckman Coulter). The generated libraries were amplified using the KAPA HiFi Hotstart Ready Mix and Nextera Index Kit 1 (i7) and 2 (i5) primers (Illumina). Amplified libraries were purified using a 0.65x AMPure XP beads (Beckman Coulter) followed by a 1.8x AMPure XP beads (Beckman Coulter) purification. Library concentration was measured using the dsDNA Fluorescence Quantification Assays (DeNovix), and library size was determined using the BioAnalyzer High Sensitivity DNA Kit (Agilent Technologies). DNA libraries were sequenced with an Illumina NextSeq 500, and 38-bp paired-end reads were generated.

### Single-cell RNA-sequencing sample preparation

After harvesting, organoids were dissociated using TrypLE (Gibco). Viable single cells were sorted in 384- well PCR plates (Bio-Rad) using the Becton Dickinson Aria (BD Biosciences) sorter with a 100 μm nozzle, based on forward and side scatter gating. Prior to sorting, plates were primed with oligos (5’-3’) containing the T7 promoter, a 5’ Illumina adapter, unique molecular identifiers (UMIs), a unique cell barcode and the oligo-(dT)N for tagging of individual mRNA molecules within each cell (63). Immediately after sorting, plates were centrifuged for 2 minutes at 1,200 x g and 4°C and stored at −80°C until library preparation. Libraries were prepared using a modified CEL-Seq2 protocol (64). In brief, a microdispenser machine Nanodrop II (BioNex) was used to dispend in each well the ERCC spike-in control RNAs (1:50,000, Invitrogen), reagents used for the reverse transcription (RT) reaction such as Superscript II (Invitrogen) and RNasein Plus (Promega), and for the second strand synthesis such as *E. coli* DNA polymerase and *E. coli* DNA ligase (New England Biolabs). Second strand synthesis was done in the presence of *E. coli* RNase H (Invitrogen). Double stranded cDNA from each well was pooled together and purified with AMPure XP beads (Beckman Coulter). *In vitro* transcription was performed overnight according to the Ambion MegaScript IVT kit protocol (Invitrogen), followed by an exonuclease digestion step with EXOI/rSAP-IT (Applied Biosystems). The amplified RNA was later fragmented and cleaned up using AMPure XP beads (Beckman Coulter). Samples were then subjected to library RT using random octamers (63) and to cDNA amplification thereafter using the Phusion High-Fidelity DNA Polymerase (New England Biolabs). A forward primer sequence against the 5’ Illumina adapter (63) and a reverse primer sequence against the octamers overhang incorporated during RT were added to the amplification mix. The reverse primer contained an Illumina 3’ adapter sequence and an index sequence to uniquely identify each library (NextFlex DNA barcodes, PerkinElmer). Libraries were purified and quantified using the dsDNA Fluorescence Quantification Assays (DeNovix) and fragment sizes were assessed with the BioAnalyzer High Sensitivity DNA Kit (Agilent Technologies). The libraries were sequenced on an Illumina NextSeq 500, and 63-bp paired-end reads were generated.

### Bulk RNA-sequencing analysis

Paired-end reads were aligned to the mouse mm10 genome using the STAR RNAseq-aligner v2.7.9a (65), with –sjdbOverhang set to 100. Mapped reads were counted with HTSeq-count v0.13.5 (66) and the following settings: -r pos –s reverse –t exon. Normalization and log_2_-transformation of data was done using the DESeq2 package v1.34.0 (67). Genes with an adjusted p-value < 0.05 were considered statistically significant. Expression heatmaps were created with ComplexHeatmap v2.10.0 (68).

### Bulk ATAC-sequencing analysis

Preprocessing of reads was done automatically with atac-seq workflow from the seq2science pipeline v.0.3.1 (69). In short, reads were trimmed using fastp v0.20.1 (70) and aligned to the mouse mm10 genome using bwa-mem v0.7.17 (71). Mitochondrial reads, multipmapper reads, mapped reads below a minimum mapping quality of 20 or reads inside the ENCODE blacklist (72) were filtered out. Reads were shifted +4 bp/−5 bp on the positive/negative strand to account for the *Tn5* inserting bias. Duplicate reads were removed with picard MarkDuplicates v2.21.2 (73). Peaks were called using macs2 v2.2.7 (74), with the settings --shift −100 --extsize 200 --nomodel --keep-dup 1 --buffer-size 10000 in BAM mode. Being paired-end reads, the option macs2_keep_mates was set to true.

### ChIPmentation analysis

The ChIPmentation reads were processed with the chip-seq workflow from seq2science v.0.3.1 (69), taking the 1% input samples of either control or RANKL-treated organoids as background for peak calling. The workflow of preprocessing, alignment, deduplication and quality filtering was similar to the workflow described for the ATAC-seq analysis, but without shifting sequencing reads in alignment or during peak calling.

For the ONECUT2 ChIPmentation, we used Diffbind (75) (v.3.4.11) to identify differentially bound genomic loci between RANKL- and control-treated organoids, with an FDR-cutoff of 0.05. For visualization of the results, we used the *coverage_table* function from GimmeMotifs to generate a table of log-transformed counts for these differential regions for RANKL and control samples. The top 250 peaks based on the fold change of RANKL over control were used to generate a ONECUT2 ChIP gene set.

Lastly, HNF4G ChIPmentation samples from mouse small intestinal organoids grown in EN medium and control were obtained from GSE114113 (12) and processed with the chip-seq workflow from seq2science (v.0.7.2), without input controls for peak calling. We used GREAT (76) to assign each peak to the nearest gene and subsequently defined a EN-specific HNF4G target gene set by selecting genes near peaks from EN-cultured organoids that were not present in the control-samples, and subsequently selecting the top 250 peaks based on the fold change of the peak over background to generate an HNF4G ChIP gene set.

### Integration and analysis of bulk sequencing data

ATAC-seq peaks generated for each replicate by seq2science, for both control and RANKL-treated conditions, were combined in one peak file using the *combine_peaks* script from the package GimmeMotifs v0.16.0 (77) with the window parameter (-w 4000) (generating roughly 60,000 accessible sites). ChIPmentation alignments, corresponding to the H3k27ac signal, were quantified at these regions using the *coverage_table* function from GimmeMotifs v0.16.0 (77) with a window of 2 kb. Differential H3k27ac signal (padj < 0.05) was defined with the DESeq2 package v1.34.0 (67) and used as input for the tool *gimme maelstrom* to detect differential motifs, without filtering out redundant motifs. A z-score cut-off of > 1.64 and < −1.64 was used to define enriched motifs in either control or RANKL-treated conditions. Motifs were matched to transcription factors with the *gimme motif2factors* conversion table provided by GimmeMotifs (77). Only motifs for which the corresponding transcription factor was differentially expressed (padj < 0.05) in the bulk RNA-seq data were included for visualization in an integrative heatmap, which was generated with ComplexHeatmap v2.10.0 (68).

### Gene-regulatory network analysis with ANANSE

The *ananse binding* command line tool from ANANSE v0.3.0 (44) was used to predict treatment-specific transcription factors binding profiles, using the ATAC-seq and ChIPmentation outputs from seq2science. Enhancer regions were specified with the combined peak summit file containing identified ATAC-seq peaks from both RANKL-treated and control conditions. The function *ananse network* was run to infer treatment-specific gene-regulatory networks. For this, binding profiles from *ananse binding*, gene-level transcript-per-million (TPM) data for each condition and the mm10 as the genome assembly were used. The mm10 genome assembly was downloaded from UCSC using genomepy v0.9.1 (78). To obtain a gene-level TPM file, reads in the RNA-seq FASTQ files (.fastq) were quantified using the command line tool Kallisto v0.46.2 (79). The prebuild Kallisto index constructed from the Ensemble reference mouse transcriptome GRCm38 was downloaded from https://github.com/pachterlab/kallisto-transcriptome-indices/releases and used. Using tximport v1.14.2 (80) the expression of all isoforms corresponding the same gene was summed up and identifiers were changed to gene-level. Lastly, the *ananse influence* function was used to generate a differential gene-regulatory network, based on the difference in the interaction score calculated per factor between RANKL treatment and control conditions. Number of top edges used to define the differential network was set to 500.00 (-i 500_000).

### Analysis of single-cell RNA-sequencing data

Raw FASTQ. files were demultiplexed, aligned and annotated to the mouse GRCm38 genome indexed with sequences from the ERCC ExFold RNA Spike-In Mixes (invitrogen), and counted using the kallisto| bustools wrapper kb_phython v0.26.3 (79,81). Pre-processing and quality control was performed using the Scater package v1.10.1 (82). To exclude low-quality cells from further analysis, any cell with < 500 detected features, < 1,000 total transcripts, > 40% mitochondrial transcripts or > 20% ERCC transcripts were removed from the datasets. Normalization, natural log-transformation, scaling (with “nCount” and “nFeature” as variables to regress), and clustering analysis were performed using Seurat v3.1.5 (83). We determined the 2,000 highly variable genes using the variance stabilizing transformation (VST) method. Dimensionality reduction was performed by principal component analysis (PCA), and was followed by Louvain clustering using the *FindNeighbors* function with the top 30 principal components (PCs) and the *FindClusters* function with a resolution set to 1.6 for the RANKL treatment dataset (21,099 genes across 6,636 cells), and the top 20 PCs and a resolution of 1.5 for the dataset of RANKL and CSRM617 combined treatment (9,257 genes across 1,903 cells). Data was projected in a two-dimensional space using the Uniform Manifold Approximation and Projection (UMAP) method. Label transferring from clusters 6, 12 and 17 from the RANKL treatment dataset (21,099 genes in 820 cells) to clusters 3, 9, 10 and 12 from the dataset of RANKL and CSRM617 combined treatment (9,257 genes in 427 cells) was done using the Seurat *FindTransferAnchors* function using the RANKL treatment dataset selected clusters as a reference.

### Trajectory inference

RNA velocity estimation was done using velocyto.R v.0.6 (27). The raw counts corresponding to spliced and unspliced RNA forms for each gene, generated by the kallisto| bustools wrapper and stored as separate assays in the Seurat object, were used by velocyto.R to create an RNA velocity map. The RNA velocity map was then projected as a grid with arrows onto the UMAP space that was generated by Seurat with the function *show.velocity.on.embedding.cor*.

For the pseudotime analysis, raw counts of cells from the Seurat clusters 6, 12 and 17 from the RANKL treatment dataset that passed the Scater quality assessment were processed with the R package Monocle3 v1.0.0 (84) with default settings. The *preprocess_cds* function of Monocle3 normalized by log and scaled the data. The function also calculated a low-dimensional space used as input for dimensionality reduction with a UMAP. With the *learn_graph* function, Monocle3 used a principal graph-embedding procedure based on the SimplePPT algorithm (85,86) to represent on the UMAP the possible paths cells can take as they develop. The pseudotime was calculated across these paths by setting the furthest away cells from cluster 17 as time zero. Lastly, we used *graph_test*, a function based on the Moran’s statistical test (87) to define the pseudotime-dependent genes and the *find_gene_modules* function to group these genes into modules based on their co-expression across clusters.

### Motif activity prediction with SCEPIA

Using SCEPIA v0.5.1 (https://github.com/vanheeringen-lab/scepia) a list of transcription factors of interest was established by predicting transcription factor motif activities based on the single-cell RNA-seq data. The normalized and scaled spliced RNA counts and associated annotation generated with Seurat (v3), including highly variable genes and clustering, were exported to an AnnData object in Python v3.9.6. First, H3K27ac profiles were assigned to the single cells using a H3K27Ac reference database. The ENCODE H3K27Ac ChIP-seq database of 121 human cell lines and tissues included in SCEPIA was used as a reference (88,89). The regulatory potential, a summarized, distance-weighted measure of H3K27ac within 100kb of the gene transcription start site, was calculated for every gene, according to Xu *et al*. (44) similar to Wang *et al*. (90). Using the top 2,000 highly variable genes from the scRNA-seq data, cells were matched to H3K27Ac profiles using Ridge regression with the normalized and scaled gene expression level as the response variable and the H3K27Ac regulatory potential as the explanatory variable. The output of this regression was a matrix with per single cell a coefficient for each cell type (single cell x cell type matrix). Only the top 50 scoring cell types (ordered by the mean absolute coefficient over the clusters) were kept and from these the top 10,000 highest variable enhancers, based on the genome-wide H3K27Ac data, were selected for motif activity analysis using the Bayesian Ridge method with GimmeMotifs v0.16.1 (77) resulting in a motif x cell type matrix. The predicted motif coefficients (“motif activity”) for each H3K27Ac reference cell type were combined into single cell motif activities using a dot product of the single cell x cell type matrix and the motif x cell type matrix. Pearson correlations between single cell motif activity and the single cell expression levels of the transcription factors binding the motifs, based on the default database of GimmeMotifs, were calculated, followed by 100,000 random permutations to determine a permutation-based p-value. The permutation-based p-value was combined with the p-value of the correlation using Fisher’s method and corrected for multiple testing using the Benjamini-Hochberg correction. From the list of transcription factor-motif combinations, the hits with the highest positive correlation coefficients and an FDR ≤ 0.1 were selected. The *gimme logo* tool from GimmeMotifs (77) was used to generate motif logos.

### Gene-regulatory network analysis with pySCENIC

Raw counts from the Seurat object were used as input for pySCENIC v0.11.0 (91). First, gene-regulatory networks between transcription factors and putative target genes were inferred with the *grnboost2* function from the Arboreto package v0.1.6 (92) part of pySCENIC, using a mouse transcription factor database available from the TFCat portal through pySCENIC. The *ctx* command kept only transcription factor motifs within 10 kb from the target transcription start site (TSS) and pruned indirect targets from dataset based on cis-regulatory cues. For this, the motifs database motifs-v9-nr.mgi-m0.001-o0.0, the c/s-target databases mm10__refseq-r80__500bp_up_and_100bp_down_tss.mc9nr and mm10__refseq-r80__10kb_up_and_down_tss.mc9nr were used. As a default option, only transcriptional regulatory units (regulons) with a positive correlation between the expression of a transcription factor and its targets were used for further analysis. The area under the curve (AUC) scores (regulon activity) were calculated for every cell using the *aucell* function with default options and later averaged per Seurat cluster. Data was visualized using gplots v3.1.1 (93).

### Gene-set enrichment analysis

Gene-set enrichment analysis (GSEA) was performed using the fgsea package v1.19.3 (94). Mouse *in vivo* intestinal cell-type gene sets were extracted from (9) by selecting the top 250 differentially expressed genes with an FDR (Q, max) < 0.05 and a mean natural-log fold change > 0.5. To create the ONECUT1-, ONETCUT2- and ONECUT3-specific signatures, RNA-seq data from Data ref: (48) was downloaded using the download-fastq workflow from seq2science and re-analyzed with DESeq2 package v1.34.0 (67). Top 250 differentially expressed genes (padj < 0.05), unique for each overexpression condition and with a Iog_2_ fold change > 0.5 over control were used to create the gene sets. Lastly, genes with a padj < 0.05 and a mean natural-log fold change > 0.5 were used to create the cluster 12 gene set from the RANKL treatment single-cell dataset. For all experiments, ranked files with log_2_ fold change expression values (padj < 1) were subjected to GSEA.

### Real-time semi-quantitative RT-qPCR

Mice small intestines were washed with 1X PBS to remove excess of RNAlater (Invitrogen) and 15-20 mg of tissue were lysed with 1 mL of TRIzol (Ambion) and incubated overnight at −20°C. At the next day, tissue was homogenized using RNAse-free pestles, 14G (1.6 mm) and 21G (0.8 mm) needles. Lysates were centrifuged for 5 minutes at 12,000 × g and 4°C. Supernatant was processed for RNA isolation following TRIzol manufacturer’s instructions. After this, a total of 30 μg of isolated total RNA per replicate was purified with the Quick-RNA Microprep Kit (Zymo research) and 1 μg of purified RNA was used for cDNA synthesis using the iScript cDNA Synthesis Kit (Bio-Rad). Real-time PCR analysis was performed using the iQ SYBR Green Supermix (Bio-Rad) in a CFX96 Real-Time system (Bio-Rad). Crossing-point (Cp) values were determined by the CFX Manager software version 3.0 (Bio-Rad). Expression levels of Gapdh were used for normalization. Relative gene expression levels were calculated according to Pfaffl *et al*. (95). Primers for RT-qPCR can be found in Additional file 2: Table S8.

### Flow-cytometry analysis

After harvesting, organoids were dissociated using TrypLE (Gibco). Organoid cells were counted using the Muse Cell Analyzer (Merck) and a total of 100,000 viable cells per well and in triplicates per treatment were seeded in conic v bottom 96-well plates (greiner bio-one). After washing with 1% PBA buffer (1% Bovine serum albumin (Sigma) in 1x PBS), cells were incubated with rat anti-mouse GP2 antibody (MBL International Corporation, D278-5, RRID:AB_11160946) in a 1:10 dilution in 100 μl of 1% PBA buffer per well. Incubation was done for 30 minutes at 4°C and was followed by three washing steps with 1% PBA buffer. All washing steps were done by centrifugation of 3 minutes at 200 x g and 4°C. Cells were lastly fixed with 1% formaldehyde for 20 minutes at room temperature and in dark. After removal of the formaldehyde solution and one washing step, cells were resuspended in 1% PBA before being analyzed in the MACSQuant Analyzer 10 (Miltenyi Biotec). FITC signal (LGR5^+^ cells) and PE signal (GP2^+^ cells) were detected in cell population selected based on forward and side scatter gating, and pseudocolor density plots were generated using FlowJo v10 (BD Biosciences).

### Enzyme-linked immunoassay (ELISA)

Measurement of secretory IgA (SIgA) in mouse faeces samples after treatment with the ONECUT2 inhibitor CSRM617 Hydrochloride (Sigma-Aldrich) or PBS control was done using the SIgA Mouse Uncoated ELISA kit (Invitrogen) following manufacturer’s instructions. Faecal supernatants were diluted to a final assay concentration of 10 ng/mL. A solution of 2N H2SO4 was used to stop the reaction and the concentration of SIgA was deduced from the standard curve made with the IgA standards provided by the kit.

### Live imaging and immunohistochemistry

Whole-mount immunofluorescence staining of organoids was performed in conical bottom polystyrene 12 mL tubes (Greiner). Briefly, after harvesting of organoids, these were transferred to the round-bottom 15 mL tubes pre-coated with 1% PBA. Two washing steps with ice-cold 1x PBS were carried out by gently resuspending the organoids and letting them sit at the bottom of the tubes before discarding supernatant. Organoids were then fixed in 4% formaldehyde for 15 minutes followed by two washing steps with ice-cold 1x PBS. Permeabilization of organoids was done with 0.5% Triton in 1x PBS for 30 minutes. This was followed by two washing steps and a quenching step with 20 mM Glycine in 1x PBS for 30 minutes. Unspecific signal was then blocked with 1% PBA for 30 minutes. Three washing steps with 1% PBA were done before overnight incubation with primary antibodies rat anti-GP2 (MBL International Corporation, D278-3, RRID:AB_10598188), diluted 1:100 in blocking solution. To prevent organoids from sitting at the bottom of the tube, incubation was carried out in a shaker at 45 rpm. Organoids were then washed three times with 1x PBS and incubated with Alexa Fluor 647 Goat anti-Rat IgG secondary antibody (Invitrogen, A-21247, RRID:AB_141778) diluted 1:400 in blocking solution, for 1.5 hours in a shaker at 45 rpm and covered from light. Antibodies were washed out three times with 1x PBS. Staining with methanolic phalloidin (Thermo Fisher Scientific) diluted in blocking solution was performed for 20 minutes and followed by three washing steps. Finally, nuclear counterstaining with Hoechst (50 μg/mL, brand) was done for 20 minutes. Organoids were washed three times more and mounted on 22 mm-diameter glass bottom dishes (WillCo Wells) with Ibidi mounting medium (Ibidi). For live microscopy, EDU labelling was performed using the Click-iT EdU Imaging Kits (Invitrogen) and following manufacturer’s protocol on organoids embedded in single BME domes, seeded on 8-well chamber slides (Ibidi).

Formalin-fixed paraffin-embedded mice small intestines were sectioned 4 μm-thick and mounted on Superfrost Plus slides (Thermo Fisher Scientific). Deparaffination was performed overnight at 37°C and re-hydration steps were followed by heat-induced epitope retrieval with boiling sodium citrate (pH9) for 10 minutes. Blocking of endogenous perodixase activity was done with 1% H_2_O_2_ for 15 minutes and was followed by blocking of unspecific binding with 2% PBA for 1 hour. GP2 or SPIB primary antibodies (1:100) were incubated overnight in 1% PBA. For detection of GP2 protein, sections were incubated for 30 minutes with biotinylated Rabbit IgG anti-rat (H+L) secondary antibody (Vector Laboratories, BA-4001, RRID:AB_10015300) 1:200 in 1% PBA and 30 minutes with an ABC mix solution from the VECTASTAIN Elite ABC Reagent kit (Vector Laboratories, PK-6101, RRID:AB_2336820). To reveal the reaction, sections were incubated with Bright DAB solution (Immunologic) for 5 minutes followed by 10-seconds nuclear counterstaining with Hematoxylin, dehydration and mounting with Permount mounting media (Fisher Chemical). Overnight incubation with sheep anti-SPIB antibody (R&D Systems, AF7204, RRID:AB_10995033) diluted 1:100 in blocking solution was followed by a 1-hour incubation with Alexa Fluor 568 Donkey anti-Sheep IgG (H+L) secondary antibody (Invitrogen, A-21099, RRID:AB_2535753) diluted 1:400 in blocking solution. Slides were mounted with Fluoromount-G with DAPI mounting medium (Invitrogen). Except for first antibody incubations, all steps for immunohistochemistry were done at room temperature.

### Image Analysis

Live-cell and three-dimensional images were captured with a Leica TCS SP8 microscope (Leica). For live imaging the organoids were held at 37°C. Organoids were imaged in XYZ mode using an HCX PL APO 63x/1.2 water immersion objective. Post-acquisition analysis of fluorescence signal was performed manually using ImageJ version 1.53c (96). Bright-field and fluorescence imaging of mouse tissue sections was carried out in the Axio Observer 7 microscope with Sample Finder AI (Zeiss) using a 20x/0.8 objective.

Brightfield images of organoids were fed to ImageJ and processed using the OrgM macro to measure circularity (97) defined by the formula 4π(area/perimeter^2^). For analysis thresholding mode was enable and watershed mode was disabled. Circularity was plotted for each treatment.

### Statistical analysis

Statistical difference in organoid circularity obtained by ImageJ, fluorescence signal detected by Flow cytometry, relative expression obtained by RT-qPCR or fluorescence signal corresponding to IgA levels were calculated using an unpaired t test in the Prism software version 5.03 (GraphPad). Bars graphs represent the mean ± SEM. The n numbers are provided in figure legends or in the Methods section corresponding to each technique. A p-value of < 0.05 was considered statistically significant and p < 0.05 was represented by one star (*), p < 0.01 was represented by two stars (**) and p < 0.001 was represented by three stars (***).

## Supporting information

Luna-Velez_Additional file 2_Tables

## Declarations

### Ethics approval and consent to participate

For the mouse experiments, all experimental protocols and procedures were approved by the Institutional Animal Care and Use Committee (IACUC) at Cedars-Sinai Medical Center and the Comparative Medicine Department. All relevant ethical regulations, standards, and norms were rigorously adhered to.

### Consent for publication

Not applicable.

### Availability of data and materials

All sequencing data generated in this study (*i.e*., FASTQ files) as well as processed data (*i.e*. raw and normalized counts) have been deposited at GEO and are publicly available in the SuperSeries record GSEXXXXXX which links to SubSeries for ATAC-sequencing (GSEXXXXXX), ChIP-sequencing (GSEXXXXXX), bulk RNA-sequencing (GSEXXXXXX) and single-cell RNA-sequencing data (GSEXXXXXX).

The code to carry out the analysis described in this study with SCEPIA v0.5.1 is provided as Jupyter Notebooks and available at https://github.com/Rebecza/Mcell_differentiation. Original code for SCEPIA (34) (https://github.com/vanheeringen-lab/scepia) can be found at https://github.com/vanheeringen-lab/scepia/blob/master/tutorials/scepia_tutorial.ipynb.

Any additional information required to reanalyse the data reported in this paper is available from Dr. Michiel Vermeulen upon reasonable request.

### Author contributions

M.V.L.V. and M.V. conceived the study and contributed to the experimental design. M.V.L.V., H.N., A.M. and C.Q. performed experiments. M.V.L.V., H.N., R.R.S. and Y.Q. performed data analysis. C.Q. and M.R.F. carried out mice treatment. S.J.v.H. created the computational tool SCEPIA. M.V.L.V. prepared the manuscript. H.N., R.R.S., Y.Q., C.Q., A.M., G.J.C.V., M.R.F., S.J.v.H. and M.V. revised and approved the final version of the manuscript.

## Acknowledgments

The authors would like to thank the members of the Vermeulen laboratory for valuable suggestions and troubleshooting. We thank Marijke Baltissen and Lieke Lamers from the Sequencing facility from the department of Molecular Biology from Radboud University (RU) for their help with next generation sequencing, also Rob Woestenenk and Tom van Oorschot from the Radboud Technology Center for Flow Cytometry, and Laura Wingens from the department of Molecular Developmental Biology at RU for their training and assistance with fluorescence-activated cell sorting and flow cytometry. We would like to thank Marieke Willemse from the Radboud Technology Center for Microscopy for the training and assistance with microscopy. The authors also thank the Pathology laboratory at the Cedars Sinai Medical Center and Birgitte Walgreen from the department of Rheumatology at Radboudumc for the technical support with tissue preparation and immunohistochemistry, and Yanpeng Xing from the laboratory of Michael Jung at the University of California, Los Angeles, who synthesized the CSRM617 compound for the *in vivo* studies. Lastly, the authors thank Jos Smits and the members from the van Heeringen group at RU for their support and fruitful discussions.

## Authors’ information

Information from corresponding authors, @VickyLunaVelez (M.V.L.V.), @svheeringen (S.J.v.H.), @LabVermeulen (M.V.).

## Supplementary information

### Additional file 1

**Fig. S1.**
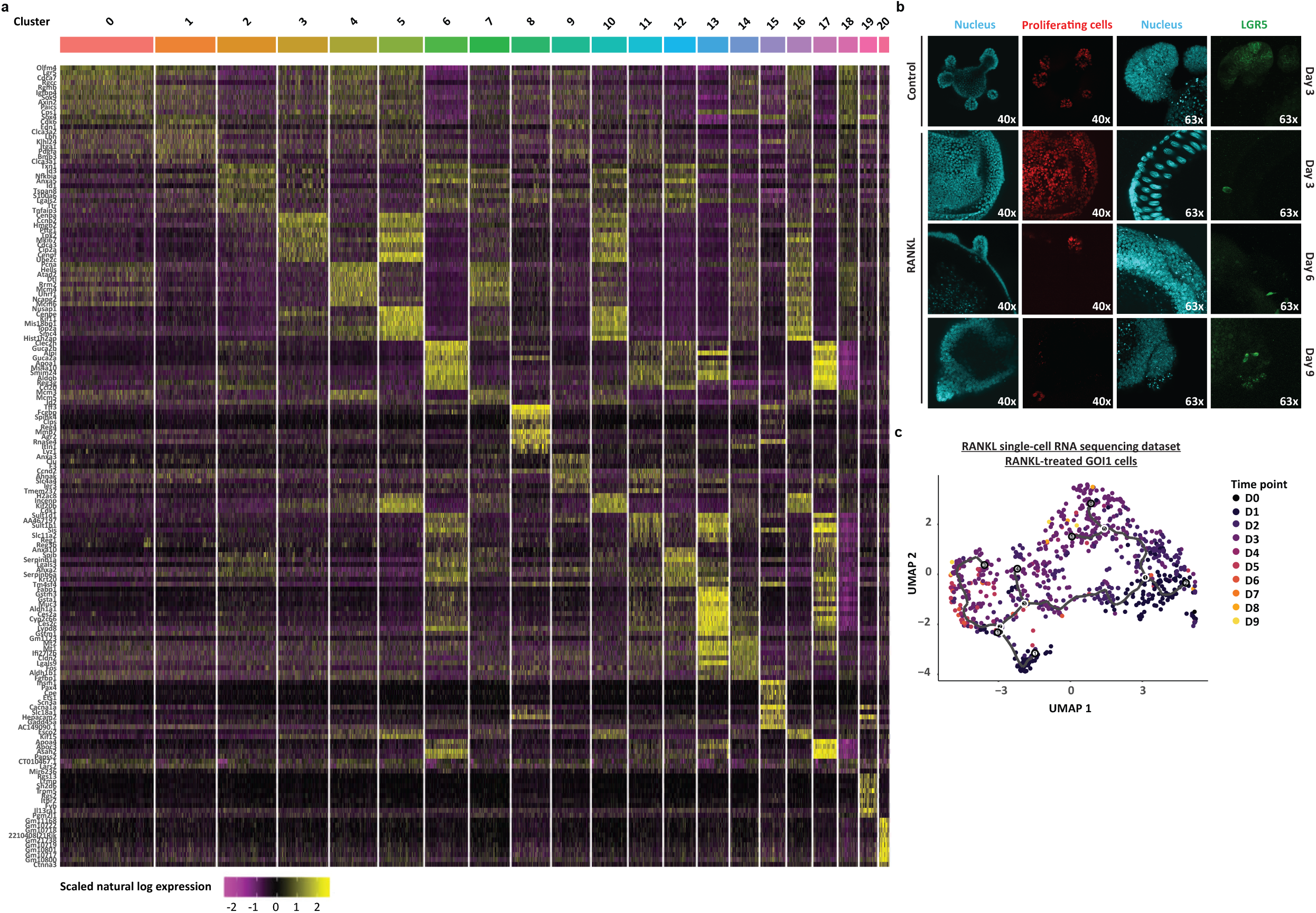
RANKL-induced transcriptional and cellular responses *in vitro*. **a** Expression heatmap of top 10 markers per cell, for each cell cluster, expressed at least in 25% of the population and with a natural-log fold change > 0.25, from the single-cell RNA-sequencing analysis of RANKL-treated organoids. **b** EdU-labelled (proliferating cells) and LGR5^+^ cells in control organoids, and in those treated with RANKL for 3, 6 and 9 days. Hoechst (nucleus) was used as counterstaining. Microscope magnification is depicted. **c** Pseudotime trajectory plot overlaid with experimental time point information of individual cells from RANKL-treated cell populations of the group of interest 1 (GOI1).

**Fig. S2.**
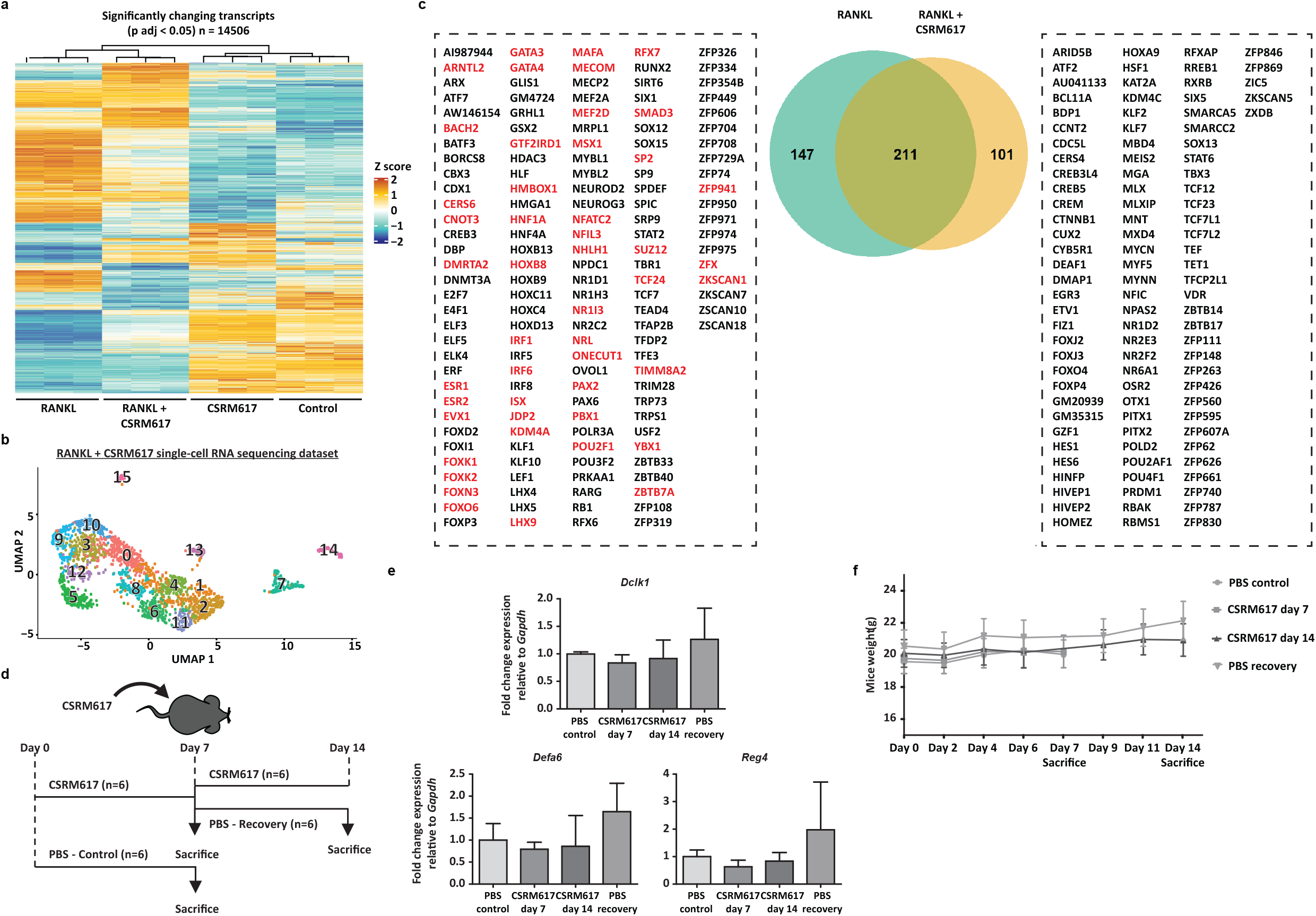
Transcriptional responses to co-treatment with RANKL and CSRM617 *in vivo*. **a** Heatmap showing the relative change in the mRNA expression of transcripts upon co-treatment with RANKL and CSRM617, compared to RANKL-alone, CSRM617 alone and control conditions. Rows show Z scores of normalized, log_2_-transformed values from significantly changing genes (padj < 0.05). b UMAP embedding of single cell transcriptome from RANKL and CSRM617 co-treatment in organoids. Each dot represents a single cell. Cell colours represent cluster identity. **c** Venn diagram with regulons in the single-cell RNA-sequencing dataset of RANKL-treated organoids and that of organoids co-treated with RANKL and CSRM617. Treatment-specific regulons are listed and ONECUT target genes are highlighted on red. **d** Work plan for *in vivo* experiments with ONECUT2 inhibitor CSRM617. Four treatment groups were designed: One group of mice treated with 50⍰mg⍰kg-1 CSRM617 for 7 days (n=6) and another group for 14 days (n=6), a recovery group consisted of mice treated with 50⍰mg⍰kg-1 CSRM617 for 7 days followed by 7 days treatment with phosphate buffered saline (PBS) and 0.2% dimethyl sulfoxide (n=6), and a control group treated with PBS and 0.2% DMSO for 7 days (n=6). **e** RT-qPCR analysis showing relative mRNA expression levels of *Dclk1, Defa6* and *Reg4* in small intestine sections from mice treated as described in **d. f** Body weight of mice treated as described in **d.**

### Additional file 2

**Table S1:** Top differentially expressed genes for each cell population (cluster) in the single-cell RNA-sequencing dataset from RANKL-treated organoids.

**Table S2:** List of transcription factor-motif combinations with the highest positive correlation coefficient between transcript expression and motif activity (FDR < 0.1) in the single-cell RNA-sequencing dataset from RANKL-treated organoids, generated by SCEPIA.

**Table S3:** List of genes constituting the KLF5, ONECUT2, TCF7L2 and HNF4G gene-regulatory networks derived from the ANANSE-generated differential gene-regulatory network of RANKL-treated organoids over control.

**Table S4:** Gene set enrichment analysis with ONECUT1-, ONECUT2- and ONECUT3-specific signatures in RANKL-treated organoids.

**Table S5:** Differential ONECUT2 binding determined by ChIPmentation and nearest genes.

**Table S6:** Composition of the regulons corresponding to the transcription factors predicted by ANANSE as drivers of *Onecut2* transcript expression.

**Table S7:** Pseudotime-dependent transcription factors within RANKL-treated cell populations of groups of interest 1 (GOI1) in the single-cell RNA-sequencing dataset from RANKL-treated organoids.

**Table S8:** RT-qPCR primers used for the detection of *Onecut2*, ONECUT2-target genes *Hnf4g, Klf5, Klf6, Hoxb8, Foxn3 and Tfcp2l1*, M cell markers *Spib, Gp2* and *Tnfaip2*, other small intestine cell types like *Dclk1, Defa6* or *Reg4*, and IgA+ B cell markers *Ccr9* and *Itgb7* in mice small intestine tissue of mice treated as in Additional file 1: Fig. S2d.

## References

1. Kraehenbuhl JP, Neutra MR. Epithelial M cells: differentiation and function. Annu Rev Cell Dev Biol. 2000;16:301–32.

2. Corr SC, Gahan CCGM, Hill C. M-cells: origin, morphology and role in mucosal immunity and microbial pathogenesis. FEMS Immunol Med Microbiol. 2008 Jan;52(1):2–12.

3. Amerongen HM, Weltzin R, Farnet CM, Michetti P, Haseltine WA, Neutra MR. Transepithelial transport of HIV-1 by intestinal M cells: a mechanism for transmission of AIDS. J Acquir Immune Defic Syndr. 1991;4(8):760–5.

4. Amerongen HM, Weltzin R, Mack JA, Winner LS 3rd, Michetti P, Apter FM, et al. M cell-mediated antigen transport and monoclonal IgA antibodies for mucosal immune protection. Ann N Y Acad Sci. 1992;664:18–26.

5. Neutra MR, Frey A, Kraehenbuhl JP. Epithelial M cells: gateways for mucosal infection and immunization. Cell. 1996 Aug;86(3):345–8.

6. Gonzalez-Hernandez MB, Liu T, Payne HC, Stencel-Baerenwald JE, Ikizler M, Yagita H, et al. Efficient norovirus and reovirus replication in the mouse intestine requires microfold (M) cells. J Virol. 2014 Jun;88(12):6934–43.

7. Fagarasan S, Honjo T. Regulation of IgA synthesis at mucosal surfaces. Curr Opin Immunol. 2004 Jun;16(3):277–83.

8. de Lau W, Kujala P, Schneeberger K, Middendorp S, Li VSW, Barker N, et al. Peyer’s patch M cells derived from Lgr5(+) stem cells require SpiB and are induced by RankL in cultured “miniguts”. Mol Cell Biol. 2012 Sep;32(18):3639–47.

9. Haber AL, Biton M, Rogel N, Herbst RH, Shekhar K, Smillie C, et al. A single-cell survey of the small intestinal epithelium. Nature. 2017 Nov;551(7680):333–9.

10. Hase K, Ohshima S, Kawano K, Hashimoto N, Matsumoto K, Saito H, et al. Distinct gene expression profiles characterize cellular phenotypes of follicle-associated epithelium and M cells. DNA Res an Int J rapid Publ reports genes genomes. 2005;12(2):127–37.

11. Sato T, Vries RG, Snippert HJ, van de Wetering M, Barker N, Stange DE, et al. Single Lgr5 stem cells build crypt-villus structures in vitro without a mesenchymal niche. Nature. 2009 May;459(7244):262–5.

12. Lindeboom RG, van Voorthuijsen L, Oost KC, Rodríguez-Colman MJ, Luna-Velez M V, Furlan C, et al. Integrative multi-omics analysis of intestinal organoid differentiation. Mol Syst Biol. 2018 Jun;14(6):e8227.

13. Wester RA, van Voorthuijsen L, Neikes HK, Dijkstra JJ, Lamers LA, Frölich S, et al. Retinoic acid signaling drives differentiation toward the absorptive lineage in colorectal cancer. iScience. 2021 Dec;24(12):103444.

14. Taylor RT, Patel SR, Lin E, Butler BR, Lake JG, Newberry RD, et al. Lymphotoxin-independent expression of TNF-related activation-induced cytokine by stromal cells in cryptopatches, isolated lymphoid follicles, and Peyer’s patches. J Immunol. 2007 May;178(9):5659–67.

15. Knoop KA, Kumar N, Butler BR, Sakthivel SK, Taylor RT, Nochi T, et al. RANKL is necessary and sufficient to initiate development of antigen-sampling M cells in the intestinal epithelium. J Immunol. 2009 Nov;183(9):5738–47.

16. Nagashima K, Sawa S, Nitta T, Tsutsumi M, Okamura T, Penninger JM, et al. Identification of subepithelial mesenchymal cells that induce IgA and diversify gut microbiota. Nat Immunol. 2017 Jun;18(6):675–82.

17. Nagashima K, Sawa S, Nitta T, Prados A, Koliaraki V, Kollias G, et al. Targeted deletion of RANKL in M cell inducer cells by the Co16a1-Cre driver. Biochem Biophys Res Commun. 2017 Nov;493(1):437–43.

18. Kobayashi A, Donaldson DS, Kanaya T, Fukuda S, Baillie JK, Freeman TC, et al. Identification of novel genes selectively expressed in the follicle-associated epithelium from the meta-analysis of transcriptomics data from multiple mouse cell and tissue populations. DNA Res an Int J rapid Publ reports genes genomes. 2012 Oct;19(5):407–22.

19. Tian H, Biehs B, Warming S, Leong KG, Rangell L, Klein OD, et al. A reserve stem cell population in small intestine renders Lgr5-positive cells dispensable. Nature. 2011 Sep;478(7368):255–9.

20. Barker N, van Es JH, Kuipers J, Kujala P, van den Born M, Cozijnsen M, et al. Identification of stem cells in small intestine and colon by marker gene Lgr5. Nature. 2007 Oct;449(7165):1003–7.

21. Boonekamp KE, Dayton TL, Clevers H. Intestinal organoids as tools for enriching and studying specific and rare cell types: advances and future directions. J Mol Cell Biol. 2020 Aug;12(8):562–8.

22. Kobayashi N, Takahashi D, Takano S, Kimura S, Hase K. The Roles of Peyer’s Patches and Microfold Cells in the Gut Immune System: Relevance to Autoimmune Diseases. Front Immunol. 2019;10:2345.

23. Kanaya T, Hase K, Takahashi D, Fukuda S, Hoshino K, Sasaki I, et al. The Ets transcription factor Spi-B is essential for the differentiation of intestinal microfold cells. Nat Immunol. 2012 Jun;13(8):729–36.

24. Kimura S, Yamakami-Kimura M, Obata Y, Hase K, Kitamura H, Ohno H, et al. Visualization of the entire differentiation process of murine M cells: suppression of their maturation in cecal patches. Mucosal Immunol. 2015 May;8(3):650–60.

25. Kimura S, Kobayashi N, Nakamura Y, Kanaya T, Takahashi D, Fujiki R, et al. Sox8 is essential for M cell maturation to accelerate IgA response at the early stage after weaning in mice. J Exp Med. 2019 Apr;216(4):831–46.

26. Creyghton MP, Cheng AW, Welstead GG, Kooistra T, Carey BW, Steine EJ, et al. Histone H3K27ac separates active from poised enhancers and predicts developmental state. Proc Natl Acad Sci U S A. 2010 Dec;107(50):21931–6.

27. La Manno G, Soldatov R, Zeisel A, Braun E, Hochgerner H, Petukhov V, et al. RNA velocity of single cells. Nature. 2018 Aug;560(7719):494–8.

28. Potten CS. Stem cells in gastrointestinal epithelium: numbers, characteristics and death. Philos Trans R Soc London Ser B, Biol Sci. 1998 Jun;353(1370):821–30.

29. Aibar S, González-Blas CB, Moerman T, Huynh-Thu VA, Imrichova H, Hulselmans G, et al. SCENIC: single-cell regulatory network inference and clustering. Nat Methods. 2017 Nov;14(11):1083–6.

30. Chen L, Toke NH, Luo S, Vasoya RP, Fullem RL, Parthasarathy A, et al. A reinforcing HNF4-SMAD4 feed-forward module stabilizes enterocyte identity. Nat Genet. 2019 May;51(5):777–85.

31. Liu H, Tang X, Srivastava A, Pécot T, Daniel P, Hemmelgarn B, et al. Redeployment of Myc and E2f1-3 drives Rb-deficient cell cycles. Nat Cell Biol. 2015 Aug;17(8):1036–48.

32. Guillermin O, Angelis N, Sidor CM, Ridgway R, Baulies A, Kucharska A, et al. Wnt and Src signals converge on YAP-TEAD to drive intestinal regeneration. EMBO J. 2021 Jul;40(13):e105770.

33. Li Q, Sun Y, Jarugumilli GK, Liu S, Dang K, Cotton JL, et al. Lats1/2 Sustain Intestinal Stem Cells and Wnt Activation through TEAD-Dependent and Independent Transcription. Cell Stem Cell. 2020 May;26(5):675–692.e8.

34. van Heeringen SJ. vanheeringen-lab/scepia: Version 0.5.1 [Internet]. 2021 [cited 2022 Jan 14], Available from: https://doi.org/10.5281/zenodo.4892888#.YeHaJLW6bs8.mendeley

35. Kim C-K, He P, Bialkowska AB, Yang VW. SP and KLF Transcription Factors in Digestive Physiology and Diseases. Gastroenterology. 2017 Jun;152(8):1845–75.

36. Nandan MO, Yang VW. The role of Krüppel-like factors in the reprogramming of somatic cells to induced pluripotent stem cells. Histol Histopathol. 2009 Oct;24(10):1343–55.

37. Nandan MO, Ghaleb AM, Bialkowska AB, Yang VW. Krüppel-like factor 5 is essential for proliferation and survival of mouse intestinal epithelial stem cells. Stem Cell Res. 2015 Jan;14(1):10–9.

38. Nigmatullina L, Norkin M, Dzama MM, Messner B, Sayols S, Soshnikova N. Id2 controls specification of Lgr5(+) intestinal stem cell progenitors during gut development. EMBO J. 2017 Apr;36(7):869–85.

39. Gregorieff A, Pinto D, Begthel H, Destrée O, Kielman M, Clevers H. Expression pattern of Wnt signaling components in the adult intestine. Gastroenterology. 2005 Aug;129(2):626–38.

40. McConnell BB, Yang VW. Mammalian Krüppel-like factors in health and diseases. Physiol Rev. 2010 Oct;90(4):1337–81.

41. Dusing MR, Maier EA, Aronow BJ, Wiginton DA. Onecut-2 knockout mice fail to thrive during early postnatal period and have altered patterns of gene expression in small intestine. Physiol Genomics. 2010 Jun;42(1):115–25.

42. Korinek V, Barker N, Moerer P, van Donselaar E, Huls G, Peters PJ, et al. Depletion of epithelial stem-cell compartments in the small intestine of mice lacking Tcf-4. Nat Genet. 1998 Aug;19(4):379–83.

43. Chen L, Toke NH, Luo S, Vasoya RP, Aita R, Parthasarathy A, et al. HNF4 factors control chromatin accessibility and are redundantly required for maturation of the fetal intestine. Development. 2019 Aug;146(19).

44. Xu Q, Georgiou G, Frölich S, van der Sande M, Veenstra GJC, Zhou H, et al. ANANSE: an enhancer network-based computational approach for predicting key transcription factors in cell fate determination. Nucleic Acids Res. 2021 Aug;49(14):7966–85.

45. Lemaigre FP, Durviaux SM, Truong O, Lannoy VJ, Hsuan JJ, Rousseau GG. Hepatocyte nuclear factor 6, a transcription factor that contains a novel type of homeodomain and a single cut domain. Proc Natl Acad Sci USA. 1996 Sep;93(18):9460–4.

46. Vanhorenbeeck V, Jacquemin P, Lemaigre FP, Rousseau GG. OC-3, a novel mammalian member of the ONECUT class of transcription factors. Biochem Biophys Res Commun. 2002 Apr;292(4):848–54.

47. Kropp PA, Gannon M. Onecut transcription factors in development and disease. Trends Dev Biol. 2016;9:43–57.

48. van der Raadt J, van Gestel SHC, Nadif Kasri N, Albers CA. ONECUT transcription factors induce neuronal characteristics and remodel chromatin accessibility. Nucleic Acids Res. 2019 Jun;47(11):5587–602.

49. Rotinen M, You S, Yang J, Coetzee SG, Reis-Sobreiro M, Huang W-C, et al. ONECUT2 is a targetable master regulator of lethal prostate cancer that suppresses the androgen axis. Nat Med [Internet]. 2018/11/26. 2018 Dec;24(12):1887–98. Available from: https://www.ncbi.nlm.nih.gov/pubmed/30478421

50. Corthésy B. Multi-faceted functions of secretory IgA at mucosal surfaces. Front Immunol. 2013;4:185.

51. Martinoli C, Chiavelli A, Rescigno M. Entry route of Salmonella typhimurium directs the type of induced immune response. Immunity. 2007 Dec;27(6):975–84.

52. Hashizume T, Togawa A, Nochi T, Igarashi O, Kweon M-N, Kiyono H, et al. Peyer’s patches are required for intestinal immunoglobulin A responses to Salmonella spp. Infect Immun. 2008 Mar;76(3):927–34.

53. Wilmore JR, Gaudette BT, Gómez Atria D, Rosenthal RL, Reiser SK, Meng W, et al. IgA Plasma Cells Are Long-Lived Residents of Gut and Bone Marrow That Express Isotype-and Tissue-Specific Gene Expression Patterns. Front Immunol. 2021;12:791095.

54. Kucharzik T, Lügering N, Rautenberg K, Lügering A, Schmidt MA, Stoll R, et al. Role of M cells in intestinal barrier function. Ann N Y Acad Sci. 2000;915:171–83.

55. Tetreault M-P, Alrabaa R, McGeehan M, Katz JP. Krüppel-like factor 5 protects against murine colitis and activates JAK-STAT signaling in vivo. PLoS One. 2012;7(5):e38338.

56. Tahoun A, Mahajan S, Paxton E, Malterer G, Donaldson DS, Wang D, et al. Salmonella transforms follicle-associated epithelial cells into M cells to promote intestinal invasion. Cell Host Microbe. 2012 Nov;12(5):645–56.

57. Golovkina T V, Shlomchik M, Hannum L, Chervonsky A. Organogenic role of B lymphocytes in mucosal immunity. Science. 1999 Dec;286(5446):1965–8.

58. Kernéis S, Bogdanova A, Kraehenbuhl JP, Pringault E. Conversion by Peyer’s patch lymphocytes of human enterocytes into M cells that transport bacteria. Science. 1997 Aug;277(5328):949–52.

59. Moolenbeek C, Ruitenberg EJ. The “Swiss roll”: a simple technique for histological studies of the rodent intestine. Lab Anim. 1981 Jan;15(1):57–9.

60. Buenrostro JD, Giresi PG, Zaba LC, Chang HY, Greenleaf WJ. Transposition of native chromatin for fast and sensitive epigenomic profiling of open chromatin, DNA-binding proteins and nucleosome position. Nat Methods. 2013 Dec;10(12):1213–8.

61. Corces MR, Trevino AE, Hamilton EG, Greenside PG, Sinnott-Armstrong NA, Vesuna S, et al. An improved ATAC-seq protocol reduces background and enables interrogation of frozen tissues. Nat Methods. 2017 Oct;14(10):959–62.

62. Schmidl C, Rendeiro AF, Sheffield NC, Bock C. ChIPmentation: fast, robust, low-input ChIP-seq for histones and transcription factors. Nat Methods. 2015 Oct;12(10):963–5.

63. Gerlach JP, van Buggenum JAG, Tanis SEJ, Hogeweg M, Heuts BMH, Muraro MJ, et al. Combined quantification of intracellular (phospho-)proteins and transcriptomics from fixed single cells. Sci Rep. 2019 Feb;9(1):1469.

64. Hashimshony T, Senderovich N, Avital G, Klochendler A, de Leeuw Y, Anavy L, et al. CEL-Seq2: sensitive highly-multiplexed single-cell RNA-Seq. Genome Biol. 2016 Apr;17:77.

65. Dobin A, Davis CA, Schlesinger F, Drenkow J, Zaleski C, Jha S, et al. STAR: ultrafast universal RNA-seq aligner. Bioinformatics. 2013 Jan;29(1):15–21.

66. Anders S, Pyl PT, Huber W. HTSeq--a Python framework to work with high-throughput sequencing data. Bioinformatics. 2015 Jan;31(2):166–9.

67. Love MI, Huber W, Anders S. Moderated estimation of fold change and dispersion for RNA-seq data with DESeq2. Genome Biol. 2014;15(12):550.

68. Gu Z, Eils R, Schlesner M. Complex heatmaps reveal patterns and correlations in multidimensional genomic data. Bioinformatics. 2016 Sep;32(18):2847–9.

69. van der Sande M, Frölich S, Smits J, Heeringen S van. seq2science [Internet]. 2020 [cited 2022 Jan 14], Available from: https://doi.org/10.5281/zenodo.4246844#.YeHmzee5OAw.mendeley

70. Chen S, Zhou Y, Chen Y, Gu J. fastp: an ultra-fast all-in-one FASTQ preprocessor. Bioinformatics. 2018 Sep;34(17):i884–90.

71. Li H. Aligning sequence reads, clone sequences and assembly contigs with BWA-MEM. 2013 Mar 16 [cited 2022 Jan 14]; Available from: http://arxiv.org/abs/1303.3997

72. Amemiya HM, Kundaje A, Boyle AP. The ENCODE Blacklist: Identification of Problematic Regions of the Genome. Sci Rep. 2019 Jun;9(1):9354.

73. Picard. Picard Toolkit. Broad Institute, GitHub Repository. [Internet]. 2019. Available from: https://broadinstitute.github.io/picard/

74. Zhang Y, Liu T, Meyer CA, Eeckhoute J, Johnson DS, Bernstein BE, et al. Model-based analysis of ChIP-Seq (MACS). Genome Biol. 2008;9(9):R137.

75. Ross-Innes CS, Stark R, Teschendorff AE, Holmes KA, Ali HR, Dunning MJ, et al. Differential oestrogen receptor binding is associated with clinical outcome in breast cancer. Nature. 2012 Jan;481(7381):389–93.

76. McLean CY, Bristor D, Hiller M, Clarke SL, Schaar BT, Lowe CB, et al. GREAT improves functional interpretation of cis-regulatory regions. Nat Biotechnol. 2010 May;28(5):495–501.

77. Bruse N, Heeringen SJ van. GimmeMotifs: an analysis framework for transcription factor motif analysis. bioRxiv [Internet]. 2018 Jan 1;474403. Available from: http://biorxiv.org/content/early/2018/11/20/474403.abstract

78. van Heeringen SJ. genomepy: download genomes the easy way [Internet]. 2017 [cited 2022 Jan 14]. Available from: https://doi.org/10.5281/zenodo.831969#.YeHrtaT9-t8.mendeley

79. Bray NL, Pimentel H, Melsted P, Pachter L. Near-optimal probabilistic RNA-seq quantification. Nat Biotechnol. 2016 May;34(5):525–7.

80. Soneson C, Love MI, Robinson MD. Differential analyses for RNA-seq: transcript-level estimates improve gene-level inferences. F1000Research. 2015;4:1521.

81. Melsted P, Booeshaghi AS, Liu L, Gao F, Lu L, Min KH (Joseph), et al. Modular, efficient and constant-memory single-cell RNA-seq preprocessing. Nat Biotechnol [Internet]. 2021;39(7):813–8. Available from: https://doi.org/10.1038/s41587-021-00870-2

82. McCarthy DJ, Campbell KR, Lun ATL, Wills QF. Scater: pre-processing, quality control, normalization and visualization of single-cell RNA-seq data in R. Bioinformatics. 2017 Apr;33(8):1179–86.

83. Stuart T, Butler A, Hoffman P, Hafemeister C, Papalexi E, Mauck WM 3rd, et al. Comprehensive Integration of Single-Cell Data. Cell. 2019 Jun;177(7):1888–1902.e21.

84. Cao J, Spielmann M, Qiu X, Huang X, Ibrahim DM, Hill AJ, et al. The single-cell transcriptional landscape of mammalian organogenesis. Nature. 2019 Feb;566(7745):496–502.

85. Mao Q, Yang L, Wang L, Goodison S, Sun Y. SimplePPT: A Simple Principal Tree Algorithm. In: Proceedings of the 2015 SIAM International Conference on Data Mining (SDM) [Internet]. Society for Industrial and Applied Mathematics; 2015. p. 792–800. (Proceedings). Available from: https://doi.org/10.1137/1.9781611974010.89

86. Mao Q, Wang L, Tsang IW, Sun Y. Principal Graph and Structure Learning Based on Reversed Graph Embedding. IEEE Trans Pattern Anal Mach Intell. 2017 Nov;39(11):2227–41.

87. Moran PAP. Notes on continuous stochastic phenomena. Biometrika. 1950 Jun;37(1-2):17–23.

88. ENCODE Project Consortium. An integrated encyclopedia of DNA elements in the human genome. Nature. 2012 Sep;489(7414):57–74.

89. Davis CA, Hitz BC, Sloan CA, Chan ET, Davidson JM, Gabdank I, et al. The Encyclopedia of DNA elements (ENCODE): data portal update. Nucleic Acids Res. 2018 Jan;46(D1):D794–801.

90. Wang S, Zang C, Xiao T, Fan J, Mei S, Qin Q, et al. Modeling cis-regulation with a compendium of genome-wide histone H3K27ac profiles. Genome Res. 2016 Oct;26(10):1417–29.

91. van de Sande B, Flerin C, Davie K, De Waegeneer M, Hulselmans G, Aibar S, et al. A scalable SCENIC workflow for single-cell gene regulatory network analysis. Nat Protoc. 2020 Jul;15(7):2247–76.

92. Moerman T, Aibar Santos S, Bravo González-Blas C, Simm J, Moreau Y, Aerts J, et al. GRNBoost2 and Arboreto: efficient and scalable inference of gene regulatory networks. Bioinformatics. 2019 Jun;35(12):2159–61.

93. Warnes G, Bolker B, Bonebakker L, Gentleman R, Huber W, Liaw A, et al. gplots: Various R programming tools for plotting data. Vol. 2, R package version. 2005.

94. Korotkevich G, Sukhov V, Budin N, Shpak B, Artyomov MN, Sergushichev A. Fast gene set enrichment analysis. bioRxiv [Internet]. 2021 Jan 1;60012. Available from: http://biorxiv.org/content/early/2021/02/01/060012.abstract

95. Pfaffl MW. A new mathematical model for relative quantification in real-time RT-PCR. Nucleic Acids Res. 2001 May;29(9):e45.

96. Schneider CA, Rasband WS, Eliceiri KW. NIH Image to ImageJ: 25 years of image analysis. Nat Methods. 2012 Jul;9(7):671–5.

97. Cai E, Thomas RA. OrgM: A Fiji macro for automated measurement of object area, diameter and roundness from bright field images. 2019; Available from: https://github.com/neuroeddu/OrgM

